# Disruption of NUAK2-Mediated RNA Splicing by Trilaciclib Reveals a Therapeutic Vulnerability in AR-Indifferent Prostate Cancer

**DOI:** 10.1101/2025.11.12.687734

**Authors:** Umar Mehraj, Uran Maimekov, Shaista Manzoor, Emily Cordova, Manasiben Patel, Ikeer Y. Mancera-Ortiz, Stefanie Howell, Zachary W. Davis-Gilbert, Mu-En Wang, Ming Chen, Jung Wook Park, Yuzhuo Wang, Jasmine Plummer, Andrew J. Armstrong, Jiaoti Huang, David H. Drewry, Antonina Mitrofanova, Everardo Macias

## Abstract

Prostate cancer remains a leading cause of cancer-related mortality in men, with aggressive, treatment-emergent androgen receptor (AR)-indifferent subtypes, including double-negative prostate cancer (DNPC) and neuroendocrine prostate cancer (NEPC), posing major clinical challenges due to limited therapeutic options. NUAK family kinase 2 (NUAK2), an AMPK-related kinase, has been implicated in tumor growth and metastatic progression; however, its functional significance and therapeutic potential in advanced prostate cancer remain largely unexplored. Here, we identify NUAK2 as a therapeutically actionable kinase dependency in AR-indifferent prostate cancer. Transcriptomic analyses across independent patient cohorts demonstrated progressive upregulation of *NUAK2* with disease progression, with the highest expression in NEPC. Immunohistochemical analysis of clinical specimens further confirmed elevated NUAK2 protein expression in NEPC relative to prostate adenocarcinoma. Genetic loss- and gain-of-function studies established NUAK2 as a functional dependency that promotes tumor cell proliferation, clonogenic growth, and tumor growth in vivo. Mechanistically, integrated proteomic analyses revealed that NUAK2 associates with spliceosomal and RNA-processing machinery, while NUAK2 perturbation induced widespread alterations in pre-mRNA splicing programs involving genes linked to mitotic regulation and oncogenic signaling. Pharmacologic studies identified trilaciclib (G1T-28), a clinically approved CDK4/6 inhibitor, as a functionally relevant NUAK2 inhibitor that directly engages NUAK2, suppresses tumor growth, and enhances the efficacy of platinum-based chemotherapy across multiple preclinical models. Collectively, these findings uncover NUAK2 as a previously unrecognized regulator of RNA splicing and therapeutic vulnerability in AR-indifferent prostate cancer and provide a rationale for repurposing G1T-28 and developing NUAK2-directed therapeutic strategies for aggressive, treatment-refractory prostate cancer.

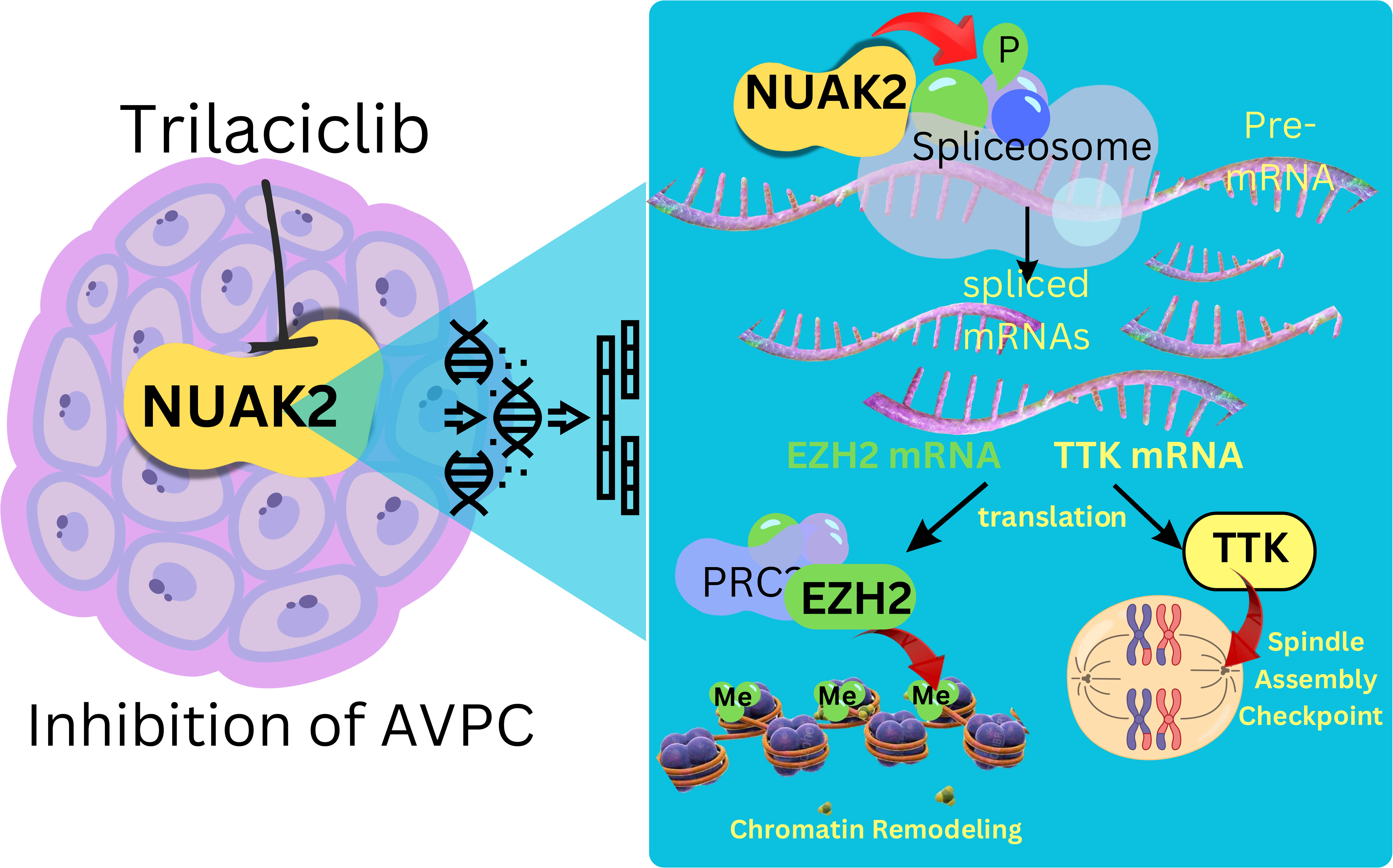

## Introduction

Prostate cancer (PCa) remains a leading cause of cancer-related mortality among men worldwide. While androgen receptor (AR) pathway inhibitors such as abiraterone and enzalutamide have significantly improved outcomes for patients with metastatic castration-resistant prostate cancer (mCRPC), the expanded use of these therapies has been accompanied by the emergence of aggressive therapy-resistant disease states (1, 2). A substantial proportion of tumors progress to AR-indifferent PCa, including DNPC and NEPC, marked by rapid progression, therapeutic resistance, and poor clinical outcomes (3, 4). Meta-analyses indicate that 9%–24% of patients receiving AR-pathway inhibitors progress to NEPC, with a median survival under one year, while rapid-autopsy cohorts show that up to 25% of prostate cancer deaths involve tumors with neuroendocrine or AR-negative phenotypes (5–7).

Genomic studies have identified recurrent alterations in AR-indifferent PCa, particularly in NEPC, which commonly harbors concurrent loss of *PTEN* and *RB1* together with *TP53* mutations (8–12). These tumors frequently exhibit amplification of lineage-defining oncogenic drivers such as MYCN and AURKA and activation of transcriptional programs that promote neuroendocrine differentiation and aggressive tumor behavior (11–13). Despite these defining genomic features, such alterations do not fully explain the profound transcriptional reprogramming and phenotypic plasticity that accompany transition to AR-indifferent states. Increasing evidence implicates dysregulated RNA processing and aberrant splicing programs as key contributors to therapeutic resistance and transdifferentiation, suggesting that regulatory proteins controlling RNA metabolism may represent actionable vulnerabilities (14–16). DNPC, defined by the absence of both AR signaling and canonical neuroendocrine marker expression, shares substantial molecular overlap with NEPC, including AR pathway suppression and activation of stemness and epithelial–mesenchymal transition programs, pointing to convergent mechanisms that drive aggressive disease progression (17, 18).

NUAK family kinase 2 (NUAK2), also known as SNARK, is a member of the AMPK-related kinase family and remains relatively understudied (19–21). Previous studies have implicated NUAK2 in tumor progression across several malignancies, including melanoma, hepatocellular carcinoma, breast cancer, and glioblastoma (22–26). We previously reported that in PCa, NUAK2 expression increases during disease progression and may be associated with metastatic potential (27). However, the functional role of NUAK2 in aggressive PCa and the molecular pathways through which it contributes to tumor progression remain largely unexplored.

Here, we investigated the role of NUAK2 in AR-indifferent PCa and its potential as a therapeutic target. We show that NUAK2 expression is elevated in AR-indifferent PCa and functions as a critical tumor dependency across multiple AR-indifferent PCa models. Integrative proteomic and transcriptomic analyses revealed that NUAK2 associates with spliceosome machinery and regulates oncogenic RNA-processing programs. Pharmacologic targeting of NUAK2 suppressed tumor growth and enhanced responses to chemotherapy. Together, these findings identify NUAK2 as a previously unrecognized regulator of RNA-processing programs and a therapeutically tractable dependency in AR-indifferent PCa.

## Methods

### Clinical datasets

NUAK2 expression analysis and pseudotime enrichment across PCa clinical progression were performed on data obtained from the Prostate Cancer Atlas (28). Transcriptomic data from the Beltran et al. cohort and Labrecque et al. cohort (GSE126078) were assessed for *NUAK2* expression levels across PCa subtypes (9, 29). Transcriptomic data from the LuCaP PDX series (GSE199596), Living Tumor Laboratory (LTL) PDX models (https://www.livingtumorlab.com/searchform.php) and LTL-331 PC to NEPC transdifferentiation model (GSE59986) were also interrogated (30, 31). For mRNA analysis across PCa cell line models, the CCLE dataset was analyzed.

### Cell lines and cell culture

Cell lines were authenticated at the Duke University Cell Culture Facility through short tandem repeat (STR) profiling and screened for mycoplasma contamination using the MycoAlert™ Plus Mycoplasma Detection Kit (Lonza). NCI-H660 and PARCB-1 cells were maintained in Advanced Dulbecco’s Modified Eagle Medium (Advanced DMEM; Thermo Fisher Scientific) supplemented with L-glutamine, B-27 supplement, recombinant human EGF (10 ng/mL), and recombinant human FGF2 (10 ng/mL). PARCB-1 cells were generated and maintained as previously described (32). HEK293T and DU-145 cells were cultured in DMEM with 10% fetal bovine serum (FBS), while PC-3, 22Rv1, and LNCaP cells were cultured in RPMI-1640 medium with 10% FBS. All cultures were incubated at 37°C in a humidified atmosphere with 5% CO₂.

### Generation of stable knockdown, knockout, and overexpression cell lines

Gene knockdown, knockout, and overexpression cell line models were generated using lentiviral transduction. Lentiviral particles were generated by co-transfecting HEK293T cells with overexpression or gene-silencing constructs, the packaging plasmid psPAX2, and the envelope plasmid pMD2.G, using the jetPRIME transfection reagent (Polyplus-transfection, Illkirch, France). The viral supernatants were collected at 48 and 72 h post-transfection, passed through a 0.22 µm filter to remove debris, and then combined in equal parts with fresh complete medium containing 8 µg/mL Polybrene (Millipore Sigma). Target cells were exposed to the viral-containing medium for 14 to 16 h, after which the medium was replaced with fresh growth medium. Selection of transduced cells was performed using antibiotics according to the respective resistance markers, with puromycin at 0.5–2 µg/mL, hygromycin B at 150–200 µg/mL, or blasticidin at 10 µg/mL. Knockdown, knockout, or overexpression was confirmed by immunoblotting.

### Generation of inducible knockdown and overexpression cell lines

Inducible NUAK2 knockdown cell lines were established using the shERWOOD-UltramiR lentiviral doxycycline-inducible shRNA platform (Skyang Bio, Huntsville, AL, USA). Unless otherwise indicated, doxycycline was used at a final concentration of 1 μg/mL in vitro. For overexpression experiments, a NUAK2 cDNA construct (pLX-NUAK2) and an empty control vector (pLX-304) were obtained from Transomic Technologies. The NUAK2 kinase-dead mutant (NUAK2_K81R) was generated by site-directed mutagenesis using the QuikChange II XL Kit (Agilent Technologies, Santa Clara, CA) and is hereafter referred to as pLX-NUAK2-KD. The mutagenesis primers used are listed in Supplementary Table S1.

### Generation of inducible NUAK2 knockout cell lines

To generate doxycycline-inducible CRISPR-Cas9 systems, cells were sequentially transduced with lentiviral vectors encoding Tet-On transactivator (pLVX-Tet3G-Blast; Addgene #128061) and doxycycline-inducible Cas9 (pLV[Exp]-Puro-TRE3G>hCas9; VectorBuilder, Chicago, IL, USA). Following transduction, cells were selected with puromycin (0.5 µg/mL) and blasticidin (10 µg/mL) for 15 days. Induction of Cas9 expression was validated by immunoblotting after a 72 h induction with doxycycline. Subsequently, Cas9-expressing cells were infected with lentiviral vectors expressing non-targeting control sgRNA or NUAK2-targeting sgRNAs (VectorBuilder, Chicago, IL, USA), then selected using hygromycin (150 µg/mL). NUAK2 knockout was confirmed by immunoblotting 96 h after doxycycline treatment.

### In vitro phenotypic assays

For drug response, cells were seeded in 96-well plates in 100 µL of medium. After 24 h, cells were treated with DMSO (vehicle), 2× drug concentrations, or doxycycline (for TRE3G inducible systems) in 100 µL of fresh medium and monitored for 5–10 days. Cell confluence was monitored using the Incucyte S3 live-cell imaging system (Sartorius, Ann Arbor, MI, USA), while cell viability was assessed with the Cell Counting Kit-8 (CCK-8, Cat# K1018, Apex Bio, Houston, TX). For drug combination assays, cells were treated in a 5 × 6 concentration matrix of the indicated compounds, and cell viability was assessed by CCK-8. Drug interactions were evaluated using the SynergyFinder Plus platform for Bliss, Loewe, ZIP, and HSA synergy scores (33). Only Bliss scores are presented in the manuscript for simplicity. For 3D spheroid assays, cells were seeded at a density of 1,000 cells per well in 96-well ultra-low attachment U-bottom plates (Prime Surface, S-BIO, Hudson, NH, USA) in 100 µL of medium. After 48 h, spheroids were treated with the indicated compounds or doxycycline at 2× concentration in an additional 100 µL of medium and growth was quantified using the Incucyte Spheroid Software Module. For clonogenic assays, cells were plated in 12-well plates at a density of 500–750 cells per well and exposed to the indicated inhibitors, doxycycline, or vehicle control. Treatments were maintained for approximately two weeks, with medium replaced every three days. At endpoint, colonies were fixed with 3.7% paraformaldehyde for 15 min and stained with 0.05% (w/v) crystal violet for 30 min at room temperature. Plates were then washed three times with water, air-dried, and imaged. To quantify colony formation, the crystal violet stain was solubilized in 10% acetic acid, and absorbance was measured at 570 nm.

### Western blotting

Cells or tissues were lysed using RIPA buffer with protease inhibitor cocktail set 1 and phosphatase inhibitor cocktails 2 and 3 (MilliporeSigma, St. Louis, MO, USA). Protein concentration was determined with DC Protein Assay (Bio-Rad, Hercules, CA, USA). For immunoblot analysis, 40–80 µg of total protein was resolved by SDS-PAGE and transferred onto 0.22 µm or 0.45 µm nitrocellulose membranes. Primary antibodies were used at dilutions ranging from 1:1,000 to 1:5,000. Membranes were developed using Pierce ECL Western (#32106), SuperSignal West Pico PLUS (#34580), SuperSignal West Dura (#34075), or SuperSignal West Femto (#34096) chemiluminescent substrates (Thermo Fisher Scientific, Waltham, MA, USA). Signal detection was performed by autoradiographic film exposure, and the films were scanned. Band intensities were quantified using ImageJ software.

### Cellular Thermal Shift Assay (CETSA)

DU-145, 22Rv1, and LNCaP cells were seeded in 150 mm plates at approximately 80% confluency. The next day, cells were treated with DMSO, HTH-02-006 (25 µM), G1T-28 (6.25 µM), or G1T-38 (6.25 µM) for 2 h. CETSA was conducted as described previously with minor modifications (34). After 2 h treatment, cells were trypsinized, collected in PBS, washed twice, and resuspended in PBS containing protease inhibitor cocktail. The cell suspension was aliquoted equally into PCR tube strips and subjected to a temperature gradient on a thermocycler for 3 min. A room temperature sample served as the control. Following thermal gradient exposure, cells were frozen on dry ice for 10 min, and samples were lysed by three freeze-thaw cycles. Lysates were collected for soluble protein analysis. Immunoblotting was performed as above, and band intensities were quantified using ImageJ and normalized to GAPDH or VINCULIN and plotted to generate protein thermal melting curves.

### Multiplexed Inhibitor Bead-Mass Spectrometry (MIB/MS)

MIB/MS was performed as described previously with minor modifications (35). Briefly, PARCB-1 cell lysates (≥5 mg protein in 4.0 mL MIB lysis buffer) were treated with DMSO or G1T-28 (100 nM, 1 µM, 10 µM) for 1 h on ice. Lysates were then applied to Bio-Rad Poly-Prep Columns (Cat# 731-1550) packed with 350 µL of a 50% slurry of multiplexed inhibitor bead (MIB) matrix containing Shokat, PP58, Purvalanol B, and UNC-21474 (14% each) and VI-16832 and Ctx-0294885 (22% each) on ECH Sepharose 4B. Columns were equilibrated in high-salt MIB wash buffer (5 mM HEPES, 1 M NaCl, 0.5% Triton X-100, 1 mM EDTA, 1 mM EGTA, pH 7.5), loaded with lysates adjusted to 1 M NaCl. Columns were sequentially washed with high-salt, low-salt MIB wash buffer (5 mM HEPES, 15 mM NaCl, 0.5% Triton X-100, 1 mM EDTA, 1 mM EGTA, pH 7.5), and 0.1% SDS in low-salt MIB wash buffer. Bound kinases were eluted with SDS/Tris buffer at 95 °C, reduced (DTT), alkylated (iodoacetamide), concentrated on 10-kDa filters (Cat# UFC801008), and precipitated by methanol/chloroform extraction. Protein pellets were dried, reconstituted in 50 mM HEPES buffer, and digested overnight with sequencing-grade trypsin. Peptides were washed with ethyl acetate, desalted using C18 spin columns (Pierce Cat# 89852), dried, and resuspended in 2% acetonitrile/0.1% formic acid. LC-MS/MS analysis was performed with a Thermo Scientific Vanquish Neo UHPLC system coupled to a Thermo Scientific Orbitrap Astral Mass Spectrometer equipped with Easy-Spray Ion Source. Samples were chromatographically separated using an IonOpticks Aurora Ultimate TS series UHPLC C18 column (75 µm ID × 25 cm, 1.7 µm particle size; IonOpticks) over a 30-minute gradient. Separation was achieved with a gradient of 5–45% mobile phase B at a 300 nL/min flow rate, where mobile phase A was 0.1% formic acid in water and mobile phase B consisted of 80% acetonitrile, 0.1% formic acid. The Astral was operated in product ion scan mode for data-independent acquisition (DIA). A full MS scan (380–980 m/z) was collected on the Orbitrap at a resolution of 240,000, a maximum injection time of 5 ms, and 500% normalized AGC target. The Astral detector acquired the MS2 scans over a scan range of 150–2000 m/z with a cycle time of 0.6 s, and a 3 m/z isolation window. The normalized AGC target was set to 500% with a maximum injection time of 3 ms, and the normalized higher collision dissociation (HCD) energy was 25%. Raw data files were processed using Spectronaut (v20.0.250606.94229; Biognosys) and searched against the UniProt reviewed human database (UP000005640, containing 20,422 reviewed sequences) appended to a common cell culture contaminants database (370 entries). The following settings were used: enzyme specificity set to trypsin, up to two missed cleavages allowed, cysteine carbamidomethylation set as a fixed modification, methionine oxidation and N-terminal acetylation set as variable modifications. A false discovery rate (FDR) of 1% was used to filter all data. Imputation was disabled, and single-hit proteins were retained. Cross-run normalization was performed within Spectronaut. Kinases were identified by cross-referencing an in-house kinome database (Lee Graves Lab, UNC Chapel Hill). A minimum of two unique peptides was required for label-free quantitation (LFQ) using the LFQ intensities. Log₂ fold-change (FC) values were calculated by subtracting the log₂-transformed protein abundance in the DMSO control from that in the drug-treated sample. (See Raw and Normalized Data, Table S2).

### TiO_2_ enriched phosphoproteomics

Proteomic experiments were conducted by the Duke Metabolomics and Proteomics Core facility. NCI-H660 cells were seeded in T75 flasks at a density of 5 × 10 per flask. After 24 h, cells were treated with DMSO, HTH-02-006 (5 µM), or G1T-28 (5 µM) for 1 h, with three biological replicates per condition. Cells were pelleted, washed twice with ice-cold PBS, and lysed in 100 µL of 8 M urea in 50 mM ammonium bicarbonate. Lysates were sonicated, quantified, and 120 µg of total protein per sample was reduced, alkylated, and digested with sequencing-grade trypsin. Peptides were resuspended in 80% acetonitrile and 1% TFA, and phosphopeptides were enriched using TiO₂ tips (GL Biosciences) according to the manufacturer’s instructions. LC-MS/MS was performed on 4 µL of each sample using a Vanquish Neo UPLC system (Thermo Fisher Scientific) coupled to a Thermo Orbitrap Astral high-resolution mass spectrometer. Peptides were first trapped on a Symmetry C18 pre-column (20 mm × 180 µm; 5 µL/min, 99.9/0.1 water/acetonitrile), followed by analytical separation on a 1.5 µm EvoSep C18 column (150 µm ID × 8 cm) with a 30-minute gradient of 5–30% acetonitrile (0.1% formic acid) at 500 nL/min and 50 °C. Data were acquired in data-independent acquisition (DIA) mode with a resolution of 240,000 at m/z 200 over a scan range of m/z 380–1080 (AGC target: 4 × 10).

Raw data from LC-MS/MS runs were processed in Spectronaut (Biognosys) using a library-free approach. Files were aligned based on precursor mass and retention time. Peptide identification was performed against the Homo sapiens SwissProt database (August 2022), a common contaminant database, and reversed-sequence decoys. Search parameters included carbamidomethylation (C) as a fixed modification and oxidation (M) and phosphorylation (S/T/Y) as variable modifications. Trypsin cleavage specificity was used with precursor and fragment mass tolerances of 10 ppm and 20 ppm, respectively. Peptides were filtered at a 1% false discovery rate (FDR).

Phosphopeptide abundance was determined from MS2 fragment ion intensities. Technical reproducibility was assessed by calculating the average coefficient of variation (CV): 40.6% for SPQC and 44.3% across treatment groups. Differential phosphorylation was evaluated using log₂-transformed intensities and two-tailed heteroscedastic t-tests comparing HTH-02-006 vs. DMSO and G1T-28 vs. DMSO (Supplementary Table S3). For Gene Ontology (GO) enrichment analysis, the Metascape online platform was used (36) and enrichment plots were generated using SR plotter (37). Protein–protein interaction (PPI) network analysis was performed using interaction data from STRING (38), BioGRID (39), OmniPath (40), and InWeb_IM (41), restricted to experimentally validated interactions. Densely connected protein interaction modules were identified using the Molecular Complex Detection (MCODE) algorithm, and the resulting networks were visualized using Cytoscape v5 (42).

### NUAK2-miniTurboBioID proximity labeling

NCI-H660 cells stably expressing NUAK2-V5-miniTurbo (mTb) (Addgene #189910) or HcRed-V5-miniTurbo (mTb) (Addgene #135237) were treated with 0.5 mM biotin for 4 h, collected, washed twice with ice-cold PBS, and lysed with Pierce™ IP Lysis Buffer (Thermo Fisher) supplemented with protease and phosphatase cocktail inhibitors as aforementioned. Samples were cleared by centrifugation at 10,000 × g for 15 min and quantified. Streptavidin affinity pulldown was performed using the Pierce™ MS-Compatible Magnetic IP Kit (Thermo Fisher, Catalog No. 90408), following the manufacturer’s instructions, and samples were eluted in 50 µL of elution buffer. Eluted samples were spiked with 1 or 2 pmol bovine casein as an internal quality control. Next, samples were reduced, alkylated, and digested using sequencing-grade trypsin (Promega). Quantitative LC-MS/MS was performed using an EvoSep One UPLC coupled to a Thermo Orbitrap Astral high-resolution accurate-mass tandem mass spectrometer (Thermo). Data were acquired in DIA mode with a full MS scan from m/z 380–980 at a resolution of 240,000 at m/z 200 and a target AGC value of 4 × 10 ions. Fixed DIA windows of 4 m/z spanning m/z 380–980 were acquired in the Astral with a target AGC value of 5 × 10 and a maximum fill time of 6 ms. Following UPLC-MS/MS analyses, data were imported into Spectronaut (Biognosys) and individual LC-MS data files were aligned. For technical reproducibility, average coefficient of variation (%CV) was calculated for each protein within each unique group; 11.7% (SPQC) and 22.5% for the other two groups. Fold-change values were calculated between NUAK2-V5-mTb and HcRed-V5-mTb groups based on protein expression values and a two-tailed heteroscedastic t-test was performed on log₂-transformed data. The complete dataset, including fold-change and P values, is provided in Supplementary Table S4. For validation, samples were run on SDS-PAGE gels after streptavidin affinity pulldown and transferred onto 0.45- or 0.22-μm nitrocellulose membrane and probed for candidate NUAK2-V5-mTb biotinylated proteins. For Gene Ontology (GO) analysis, the Metascape online tool was used, and plots were generated using SR plotter (36, 37). Gene ontology networks were analyzed and visualized with Cytoscape v5 (42).

### Immunofluorescence staining

DU-145 cells were cultured on poly-L-lysine-coated coverslips for 24 h and fixed with 2% paraformaldehyde in PBS for 20 min. Cells were then permeabilized with 0.25% Triton X-100 for 15 min and blocked with 1% BSA for 30 min. Cells were incubated overnight at 4 °C with primary antibodies against NUAK2 (Novus Biologicals, NBP1-81880; 1:100) and SC35 (Sigma-Aldrich, S4045; 1:400). After washing with PBST, cells were incubated for 1.5 h with Alexa Fluor 488- or Alexa Fluor 594-conjugated secondary antibodies. Coverslips were then washed and mounted using DAPI-containing antifade mounting medium. Images were acquired using a Zeiss LSM 880 inverted confocal laser scanning microscope.

### Co-immunoprecipitation

DU-145 and NCI-H660 cells were seeded at 80% confluence in 15 cm plates or T75 flasks, respectively. Cells were washed with PBS and lysed in 1 mL of IP lysis buffer for 30 min. Cell lysates were clarified by centrifugation at 12,000 × g for 15 min at 4°C. For co-immunoprecipitations, 25 µL of Protein G Dynabeads were pre-washed and incubated in 1 mL of lysis buffer with either 5 µg mouse anti-NUAK2 antibody (Novus Biologicals, H00081788-M04) or 5 µg normal mouse IgG (Sigma-Aldrich, 12-371) at 4°C for 3 h. Clarified protein lysates were incubated with antibody-bound Dynabeads overnight at 4°C. Beads were washed three times with lysis buffer, and bound proteins were eluted with 40 µL of 3× Laemmli buffer by boiling for 5 min. Eluates were resolved by SDS–PAGE, followed by standard immunoblotting to assess candidate NUAK2-interacting proteins.

### In vitro kinase assay

To assess NUAK2 *in vitro* pharmacologic inhibition, recombinant wild-type NUAK2 (rNUAK2, 100 ng; SignalChem, #N20-10G) was incubated with G1T-28 or HTH-02-006 in kinase buffer containing DTT (1 mM) and ATP (100 µM). Recombinant kinase-dead NUAK2 (rNUAK2-K81R, SignalChem, N20-16G) was included as a negative control. NUAK2 autophosphorylation was assessed by immunoblotting with a pan-phosphoserine/threonine antibody (Abclonal, AP1475).

For NUAK2 candidate phospho-substrate assays, rNUAK2 or rNUAK2-K81R (100 ng) was incubated with rCDC40 (400 ng; Abnova, H00051362-P01) or rSF1 (400 ng; Abnova, H00007536-P01) in kinase buffer containing DTT (1 mM) and ATP (100 µM). As indicated, G1T-28 (0.5 µM) was added prior to ATP addition. All kinase reactions were performed at 30°C for 2 h and terminated by addition of 4× SDS sample buffer. Proteins were resolved by SDS-PAGE and analyzed by immunoblotting with a pan-phosphoserine/threonine antibody to assess phosphorylation. Protein loading was confirmed by immunoblotting for NUAK2, SF1, and CDC40.

### RNA sequencing and pre-mRNA splicing analysis

NCI-H660 cells were seeded in T75 flasks at a density of 5 × 10 cells per flask. The next day, cells were treated with DMSO (vehicle) or G1T-28 (5 µM) for 24 h. Cells were collected, washed twice with ice-cold PBS, and total RNA was extracted using the RNeasy Mini Kit (Qiagen) following the manufacturer’s protocol. RNA samples were subjected to DNase treatment and quality assessment prior to sequencing. High-quality RNA was processed for 150 bp paired-end RNA sequencing on an Illumina NovaSeq X Plus platform (Genewiz, Azenta Life Sciences). Raw sequencing reads underwent quality control using FastQC v0.12.1 to assess base quality scores and detect potential sequencing artifacts. Adapter sequences and low-quality bases were trimmed using AdapterRemoval v2.2.2 (43). Trimmed reads were aligned to the human reference genome (GRCh38/hg38) using STAR v2.7.11a with default parameters (44). Differential gene expression analysis was conducted using DESeq2 (45). P values were adjusted for multiple testing using the Benjamini–Hochberg method, and genes with an adjusted P ≤ 0.01 were considered significantly differentially expressed (Supplementary Table S6). Gene ontology (GO) enrichment analysis was performed using the enrichGO function from the clusterProfiler R package (v4.14.6) (46).

Major expressed transcript isoforms were identified as previously described (47, 48). Transcript abundance was quantified using Salmon (v1.10.0) (49), and for each gene, the major isoform was defined as the transcript with the highest mean transcripts per million (TPM) across all samples, provided it accounted for ≥70% of the gene’s total expression. Isoforms located on sex chromosomes (X, Y) or mitochondrial DNA (MT), as well as those overlapping other genes, were excluded. The final reference annotation comprised 8,518 major isoforms containing 87,752 exons and 65,487 introns. Splicing ratios were computed using an exon-based splice junction analysis, as previously described (47). Only exons from major isoforms with first exons >100 base pairs were included; single-exon isoforms were excluded. A ±4 bp window around the exon–intron junction (2 bp in the exon and 2 bp in the intron) was defined. Reads overlapping ≥3 bp of this window were considered unspliced, and those overlapping ≥2 bp were considered total junctional reads. Spliced reads were determined by subtracting unspliced read counts from total reads. Splicing ratios were calculated as the proportion of spliced reads relative to the total reads per junction. Read overlap was computed using the findOverlaps function from the GenomicRanges R package (v1.58.0) (50).

Multivariate Analysis of Transcript Splicing rMATS-turbo (v4.1.2) was used to quantify five major splicing events: exon skipping (SE), intron retention (RI), alternative 5′ splice sites (A5SS), alternative 3′ splice sites (A3SS), and mutually exclusive exons (MXE) (51). Representative splicing events were visualized using rmats2sashimiplot (https://github.com/Xinglab/rmats2sashimiplot), using rMATS output files (.txt) as input. The rMATS output files are provided as Supplementary Table S7. The “inc level” in sashimi plots generated by rmats2sashimiplot refers to the exon or intron inclusion level, essentially measuring how often a specific exon or intron is present in the mature mRNA, based on sequencing reads.

### Semi-quantitative RT-PCR

NCI-H660 cells were treated with either DMSO or G1T-28 (5 µM) for 24 h, while DU-145 cells were treated with DMSO or G1T-28 (2 µM) for 6 h. For NUAK2 knockdown experiments, NCI-H660 and DU-145 cells stably expressing doxycycline-inducible shRNAs (shNT, shNUAK2-1, and shNUAK2-2) were treated with doxycycline for 72 h and 60 h, respectively. Total RNA was extracted as described above, and 1 µg of total RNA was reverse transcribed using random hexamer primers and SuperScript II Reverse Transcriptase (Thermo Fisher Scientific). Semi-quantitative PCR was performed in a 20 µL reaction containing 5 ng of cDNA. Amplified products were resolved on a 1.8% agarose gel, stained with Safe DNA Gel Stain (A8743, ApexBio), and visualized using a Li-COR imaging system. Primer sequences used for splicing analysis are listed in Supplementary Table S1.

### In vivo xenograft studies

All animal experiments were performed in accordance with ethical regulations and approved protocols from the Institutional Animal Care and Use Committee (IACUC) of Duke University (Protocol: A190-23-09 and A034-25-04) and the Department of Defense Animal Care and Use Review Office (ACURO). Male NOD-SCID IL2Rγ^null (NSG) mice were used for all studies and housed in a pathogen-free facility under standard environmental conditions (65–75 °F; 40–60% humidity).

For cell line xenograft experiments, 1 × 10 cells were suspended in a 1:1 mixture of Matrigel and Advanced DMEM/F12 (Thermo Fisher) and injected into the right flank of male NSG mice. For pZip-TRE3G-ZsGreen-shRNA or TRE3G-Cas9 studies, mice bearing tumors derived from NCI-H660 or PARCB-1 cells expressing inducible constructs were switched to and maintained on doxycycline-containing (200 mg/kg) chow (Bio-Serv, Flemington, NJ, USA) once tumors became palpable (∼50 mm³). Tumor volume was measured using calipers and quantified using the formula: (length × width²)/2. Mice were euthanized once control tumors reached an approximate volume of 1,500 mm³. Tumors were excised, weighed, and a portion flash-frozen and other portions fixed in 10% neutral-buffered formalin for histopathological analysis.

For pharmacological efficacy studies, mice were randomized into treatment groups once tumors reached a palpable volume (∼50–100 mm³). HTH-02-006 was administered by intraperitoneal (i.p.) injections twice daily in a vehicle consisting of 90% 5% dextrose solution and 10% DMSO at 5 or 10 mg/kg. For G1T-28 studies, mice were randomized to vehicle (10 mM citrate buffer, pH 4), 50- or 100 mg/kg G1T-28, administered once daily by oral gavage. For patient-derived xenograft (PDX) studies, freshly excised LTL-352 tumor tissues (∼20 mg) were implanted subcutaneously into the flank of NSG male mice under sterile conditions. Once tumors reached an average volume of 40–50 mm³, mice were randomized into control or treatment groups and treated as described above with vehicle, HTH-02-006, or G1T-28.

For the orthotopic study, PC-3_Luc cells were transduced with lentiviral vectors encoding pLX-304 (empty vector), pLX-NUAK2, or pLX-NUAK2-KD and selected with blasticidin (10 µg/mL) for 7 days. Subsequently, 2.5 × 10 cells were suspended in a 1:1 mixture of Matrigel and RPMI medium (total volume 20 µL) and orthotopically implanted into the dorsal prostatic lobe of 6–8-week-old male SCID/Beige mice (Jackson Laboratory) as described previously (52). For bioluminescence imaging, mice were injected intraperitoneally with 150 µL of D-luciferin (30 mg/mL). Tumor burden was measured and quantified using the IVIS Spectrum In Vivo Imaging System (PerkinElmer, Waltham, MA).

For complete blood count (CBC) analysis, whole blood was collected into K₂EDTA-containing tubes on treatment day 26 following administration of vehicle, G1T-28, carboplatin, or the combination. CBC measurements were performed using the HESKA Element HT5 at the Duke VDL Core Facility.

### Immunohistochemistry

Tissues were fixed in 10% neutral-buffered formalin for 24 h and subsequently paraffin-embedded and sectioned into 5 μm-thick sections at the Duke Research Histology Lab core. After deparaffinization and rehydration, antigen retrieval was performed with antigen retrieval buffer (Vector Laboratories) for 15 min. The sections were washed twice followed by blocking for endogenous peroxidase (3% H2O2 in PBS) for 20 min, then blocked with 5% goat serum in TBS/0.1% BSA for 20 min at room temperature. The sections were incubated with primary antibodies: anti-NUAK2 antibody (Novus, NBP1-81880, 1:200) and anti-Ki67 (D2H10, Cell Signaling Technology, 1:400) at 4°C overnight. Next, the sections were incubated with secondary antibodies at room temperature for 1 h. After washes, the sections were incubated with ABC Elite (Vector Laboratories, Newark, CA, USA) for 30 min. The sections were washed with TBS followed by incubation with ImmPACT DAB EqV (Vector Laboratories) and deactivation with water. After incubation in 50% hematoxylin counterstain for 1 min, the samples were dehydrated and coverslipped.

### Statistical analysis

Comparisons between two groups were performed using two-sided Student’s t tests unless otherwise specified. For multiple-group comparisons, one-way or repeated measures ANOVA was applied unless otherwise specified, followed by appropriate post hoc tests. P values ≤ 0.05 were considered statistically significant. Quantification of NUAK2 expression in PCa versus NEPC or cytoplasmic versus nuclear staining in IHC was performed using H-score by a contract pathologist blinded to the study. All statistical analyses and data visualization were conducted using GraphPad Prism (version 10.6.0).

## Results

### NUAK2 is upregulated in AR-indifferent prostate cancer

We previously demonstrated that NUAK2 is essential in CRPC and that elevated NUAK2 expression is associated with increased metastatic risk following radical prostatectomy and is enriched in NEPC (27), providing a strong rationale to investigate its functional significance and therapeutic potential in aggressive AR-indifferent prostate cancer, including DNPC and NEPC. Analysis of the pooled PCa datasets in the Prostate Cancer Atlas showed a progressive increase in *NUAK2* expression from normal prostate tissue to primary PCa and metastatic castration-resistant prostate cancer (mCRPC), with the highest expression observed in NEPC (Fig. 1A) (28). Pseudotime trajectory analysis further revealed enrichment of NUAK2 expression in disease states occupying later pseudotime states, characterized by reduced AR activity and acquisition of DNPC or NEPC phenotypes. While NUAK2 expression was heterogeneous across DNPC patients, NEPC patients exhibited consistently elevated NUAK2 expression (Fig. 1A–B). Analysis of independent clinical cohorts further confirmed elevated NUAK2 expression in aggressive PCa subtypes, with significantly higher expression in NEPC than CRPC in the Beltran dataset (9) (Fig. 1C), while the Labrecque cohort (29) demonstrated heterogeneous NUAK2 expression in the DNPC cohort but consistently elevated expression in NEPC (AR⁻/NE⁺) relative to AR-positive disease states (Fig. 1D). Consistent with these findings, similarity index analysis demonstrated enrichment of *NUAK2* expression with NEPC-like transcriptional phenotype, further linking NUAK2 to neuroendocrine lineage features (Fig. 1E–F, Table S8) (9, 29).

**Figure 1.**
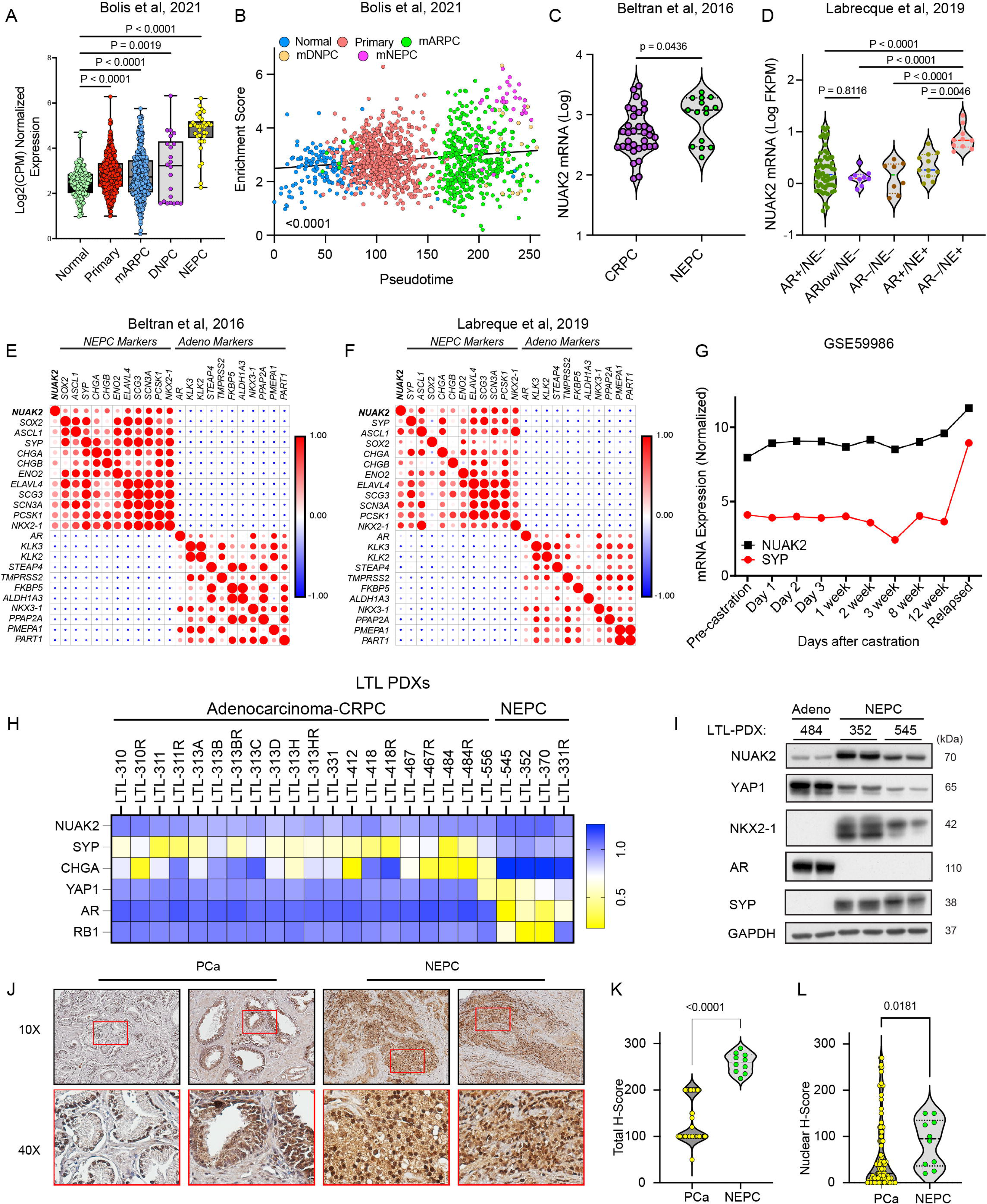
NUAK2 is upregulated in aggressive AR-indifferent PCa. (A) *NUAK2* expression across normal prostate, primary adenocarcinoma, mARPC (AR^+^), DNPC, and NEPC samples. Data from the Prostate Cancer Atlas, Bolis et al. 2021 (28). (B) Pseudotime trajectory analysis of RNA-seq data demonstrating enrichment of *NUAK2* in pseudotime states. Data from the Prostate Cancer Atlas, Bolis et al. 2021 (28). (C) Violin plots showing *NUAK2* expression in mCRPC and NEPC patient samples from the Beltran et al. dataset (9). (D) *NUAK2* mRNA expression stratified by androgen receptor (AR) and NEPC status across PCa patients in the Labrecque et al. dataset (29). (E) Similarity index analysis of *NUAK2* with neuroendocrine- and AR-related genes. Bubble color indicates the Pearson correlation coefficient (red for positive, blue for negative). Data obtained from the Beltran et al. 2016 cohort (9). (F) Similarity index analysis of *NUAK2* with neuroendocrine-related genes and AR-related genes. Bubble color indicates the Pearson correlation coefficient (red for positive, blue for negative). Data obtained from the Labrecque et al. cohort (29). (G) Longitudinal expression of *NUAK2* and *SYP* (NEPC marker) during progression from adenocarcinoma to NEPC in the LTL-331 model, illustrating concurrent induction of NUAK2 expression with neuroendocrine differentiation. Data are reproduced from the published LTL-331 longitudinal dataset (31); biological replicates were not available for statistical analysis in the original study. (H) Heatmaps illustrating mRNA expression levels of *NUAK2, SYP, CHGA, YAP1, AR,* and *RB1* across adenocarcinoma-CRPC and NEPC in the LTL PDX series. (I) Immunoblot analysis of NUAK2 and representative marker proteins of adenocarcinoma (AR, YAP1) and NEPC (NKX2-1, SYP) in LTL PDX tumor lysates. (J) Representative IHC images of NUAK2 expression in adenocarcinoma-CRPC (PCa) (n = 50) and NEPC (n = 10) tissue samples. Scale bar = 100 µm. (K) Quantification of NUAK2 protein expression from IHC images using H-score analysis. (L) Quantification of nuclear NUAK2 staining by H-score analysis in PCa versus NEPC. P values were determined by one-way ANOVA for panels (A) and (D), unpaired two-tailed Student’s *t*-test for (C, K, and L) and simple linear regression in (B). For (E) and (F), r and P values were calculated using two-tailed tests for each pairwise Pearson correlation. Corresponding r and P values for panels (E) and (F) are provided in Table S8.

To corroborate our observations from clinical specimens, we next examined NUAK2 expression across experimental models of disease progression. In PCa cell lines, NUAK2 mRNA was elevated in CRPC and NEPC models relative to adenocarcinoma cell lines, while DNPC models exhibited more heterogeneous NUAK2 expression (Fig. S1D). In the LTL-331 PDX model of adenocarcinoma-to-NEPC transdifferentiation, *NUAK2* expression was elevated in relapsed NEPC relative to preceding adenocarcinoma samples, coinciding with increased *SYP* expression (Fig. 1G) (31). Additionally, *NUAK2* mRNA expression was elevated in PDXs of NEPC histology compared with PCa adenocarcinoma in both the LTL and LuCaP PDX series (30) (Fig. 1H, Fig. S1A–C). Immunoblot analysis confirmed higher NUAK2 protein levels in NEPC PDX (LTL-352, LTL-545) tumor lysates relative to adenocarcinoma PDX (LTL-484) (Fig. 1I).

In archived PCa clinical specimens, IHC analysis demonstrated increased NUAK2 protein expression in NEPC compared with PCa adenocarcinoma tissues (Fig. 1J). Quantitative analysis confirmed significantly higher NUAK2 H-scores in NEPC specimens (Fig. 1K). In addition, NUAK2 exhibited higher nuclear H-scores in NEPC tissues (Fig. 1L).

Collectively, these findings demonstrate that NUAK2 expression increases with PCa progression and is prominently enriched in NEPC, supporting a potential role for NUAK2 in aggressive AR-indifferent disease.

### NUAK2 is required for tumor growth in AR-indifferent PCa

We next investigated the functional role of NUAK2 in tumor cell growth and survival using doxycycline-inducible NUAK2-targeting shRNAs (Fig. 2A) and TRE3G-Cas9–mediated NUAK2 knockout model systems (Fig. S2B–E). Genetic depletion of NUAK2 by shRNAs significantly suppressed cell proliferation in three cell lines (Fig. 2B–C and Fig. S2A). Consistently, TRE3G-Cas9-mediated NUAK2 knockout in NCI-H660 and PARCB-1 cells significantly reduced cell proliferation (Fig. S2D–E). In the DNPC cell line DU-145, NUAK2 depletion markedly suppressed longitudinal spheroid growth kinetics (Fig. 2D–E). Consistent with these findings, NUAK2 depletion markedly impaired clonogenic potential of DU-145 cells (Fig. 2F–G).

**Figure 2.**
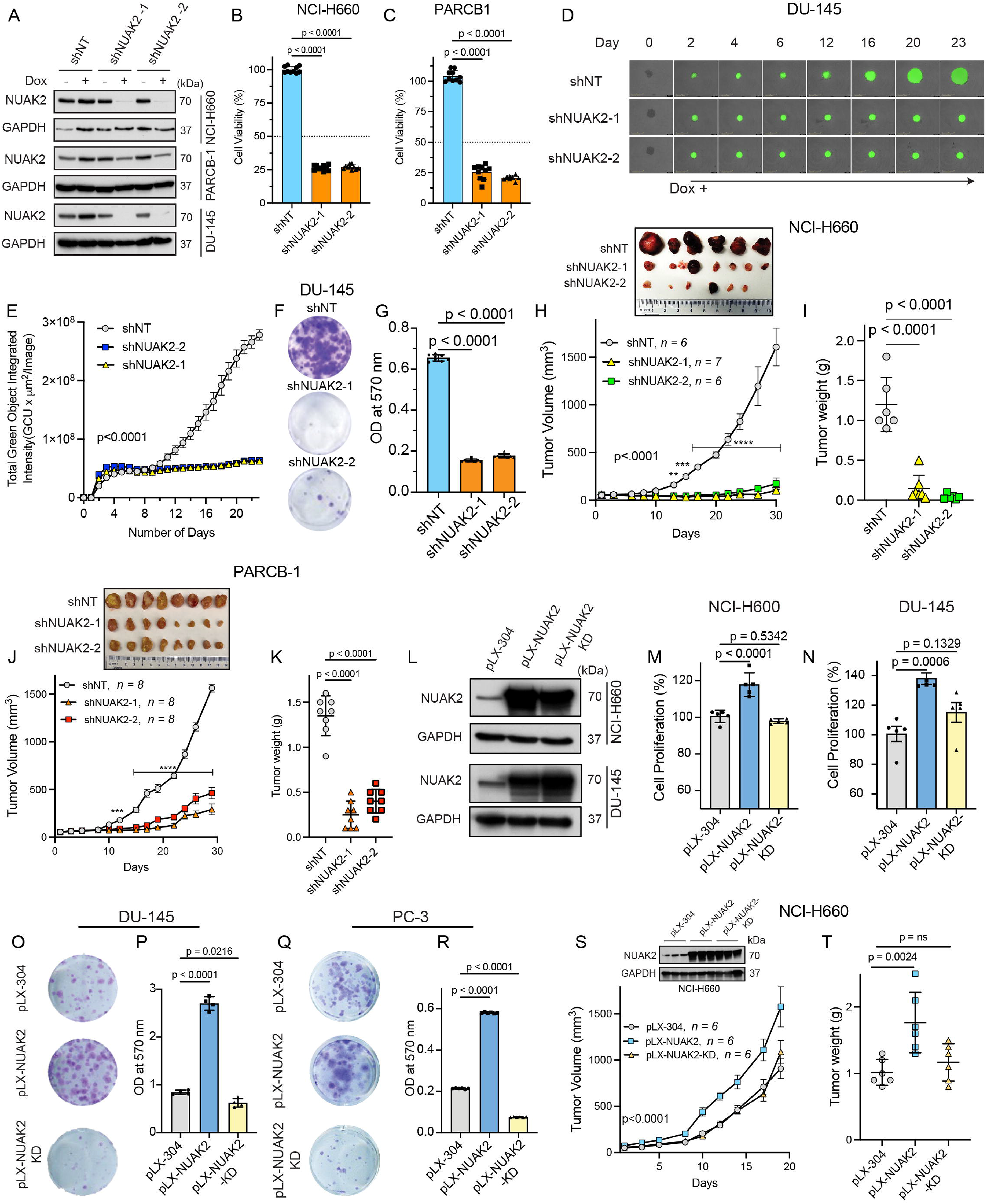
Genetic perturbation identifies NUAK2 as a critical dependency in AR-indifferent prostate cancer. (A) Representative immunoblot analysis of doxycycline-inducible NUAK2 knockdown in stable NCI-H660, PARCB-1, and DU-145 cells treated with doxycycline (1 µg/mL) for 72 h. (B) NCI-H660 and (C) PARCB-1 cell proliferation assays following doxycycline-induced NUAK2 knockdown with two independent shRNAs (shNUAK2-1 and shNUAK2-2) compared to a non-targeting control (shNT). Data are presented as mean ± SD. (D) Representative images and (E) quantification of DU-145 spheroid growth kinetics after NUAK2 knockdown versus shNT control upon doxycycline treatment. Data represent mean ± SEM, n = 6–7 biological replicates. (F) Representative images and (G) quantification of colony formation assay of DU-145 cells with either shNT or NUAK2 knockdown. Data are shown as mean ± SD from n = 10 biological replicates. (H–I) Representative endpoint images, tumor growth kinetics and endpoint tumor wet weights of NCI-H660 CDXs expressing shNT, shNUAK2-1, or shNUAK2-2. Data are means ± SEM. (J–K) Representative endpoint images, tumor growth kinetics and endpoint tumor wet weights of PARCB-1 CDXs expressing shNT, shNUAK2-1, or shNUAK2-2. Data are means ± SEM. (L) Validation immunoblots of NUAK2 expression in NCI-H660 and DU-145 cells transduced with empty vector pLX-304, pLX-NUAK2, or pLX-NUAK2-KD. (M) Proliferation assays of NCI-H660 and (N) DU-145 cells stably expressing pLX-304, pLX-NUAK2, or pLX-NUAK2-KD. Data are shown as mean ± SD. (O) Representative images and (P) quantification of colony formation assay in DU-145 cells expressing pLX-304, pLX-NUAK2, or pLX-NUAK2-KD. Data represent mean ± SD. (Q) Representative images and (R) quantification of colony formation assay in PC-3 cells expressing pLX-304, pLX-NUAK2, or pLX-NUAK2-KD. Data represent mean ± SD. (S) Representative immunoblot confirming NUAK2 expression, tumor growth kinetics, and (T) endpoint tumor weights of NCI-H660 xenografts stably expressing pLX-304, pLX-NUAK2, or pLX-NUAK2-KD. Data are presented as mean ± SEM; n = 6 mice per group. For statistical analysis, one-way ANOVA with Tukey’s multiple-comparison test was used for panels (B, C, G, I, K, M, N, P, R, and T), and two-way ANOVA with Tukey’s post hoc test was used for panels (E, H, J, and S), ****P < 0.0001, ***P < 0.001, **P < 0.01, *P < 0.05.

In both NCI-H660 and PARCB-1 in vivo xenograft models, NUAK2 shRNA knockdown markedly suppressed tumor growth and resulted in significantly reduced tumor weights at endpoint (Fig. 2H–K). Consistent with these findings, TRE3G-Cas9-mediated NUAK2 knockout in NCI-H660 xenografts significantly impaired tumor growth despite incomplete knockout efficiency, resulting in >50% reduction in tumor volume compared with controls and significantly lower tumor weights at endpoint (Fig. S2F–H).

To evaluate the requirement of NUAK2 kinase activity, we ectopically expressed empty vector (pLX-304), wild-type NUAK2 (pLX-NUAK2), or kinase-dead NUAK2 K81R (pLX-NUAK2-KD) in NCI-H660, DU-145, and PC-3 cells (Fig. 2L, Fig. S2I). Wild-type NUAK2 significantly enhanced cell proliferation (Fig. 2M–N; Fig. S2J). In contrast, NUAK2-KD largely failed to promote proliferation, although a modest increase was observed in PC-3 cells, supporting a predominantly kinase-dependent role for NUAK2 in promoting cell growth (Fig. S2J). Notably, wild-type NUAK2, but not NUAK2-KD, markedly enhanced the clonogenic capacity of DU-145 (Fig. 2O–P) and PC-3 (Fig. 2Q–R). Extending these findings *in vivo*, wild-type NUAK2 significantly accelerated NCI-H660 xenograft growth, resulting in 43% and 30% greater tumor volumes compared with the empty vector and NUAK2-KD groups, respectively (Fig. 2S). Correspondingly, endpoint tumor weights were substantially higher in the wild-type NUAK2 group than in either the empty vector or NUAK2-KD groups (Fig. 2T).

To further assess the role of NUAK2 in promoting tumor growth and metastatic dissemination, we employed the PC-3_Luc intraprostatic orthotopic xenograft model. Longitudinal bioluminescence imaging and quantitative analysis demonstrated significantly greater tumor burden in both the wild-type NUAK2 (pLX-NUAK2) and empty vector (pLX-304) groups compared with the NUAK2-KD group, whereas no significant difference was observed between wild-type NUAK2 and empty vector controls (Fig. S2K–L). However, endpoint analysis of primary prostate tumor wet weights revealed significantly higher tumor mass in the wild-type NUAK2 group relative to both the empty vector and NUAK2-KD groups (Fig. S2M). Histological analysis, including H&E staining and IHC for NUAK2 and human-specific Ki67, confirmed hepatic metastatic lesions of human tumor-cell origin (Fig. S2N). Although wild-type NUAK2 overexpression was associated with an increased number of hepatic metastatic lesions compared with controls, this difference did not reach statistical significance at humane endpoint (Fig. S2O). In contrast, expression of NUAK2-KD significantly reduced the number of hepatic metastatic lesions, as determined by quantification of human-specific Ki67-positive metastatic foci (Fig. S2N–O).

Collectively, these findings establish NUAK2 as a functional dependency in AR-indifferent PCa models, with the impaired growth and metastatic phenotypes observed upon expression of kinase-dead NUAK2 supporting a critical role for NUAK2 kinase activity.

### NUAK2 associates with spliceosome machinery and RNA-processing complexes

To elucidate the molecular mechanisms underlying NUAK2 function in NEPC, we sought to define NUAK2-associated signaling pathways and protein networks. To identify additional pharmacologic tools for interrogating NUAK2 function, we queried the Pharos database for compounds with reported NUAK2-binding activity (Fig. S3A) (20, 53). Notably, the FDA-approved CDK4/6 inhibitor trilaciclib (G1T-28) was identified as a high-affinity NUAK2-binding compound (reported pKd = 9.15). We confirmed that G1T-28 markedly suppressed NUAK2 autophosphorylation in an in vitro kinase assay (Fig. 3A). This inhibitory effect was more pronounced than that observed with HTH-02-006. These findings demonstrate that G1T-28 directly inhibits NUAK2 kinase activity in vitro and support its use as a pharmacologic tool to interrogate NUAK2 function (Fig. 3A).

**Figure 3.**
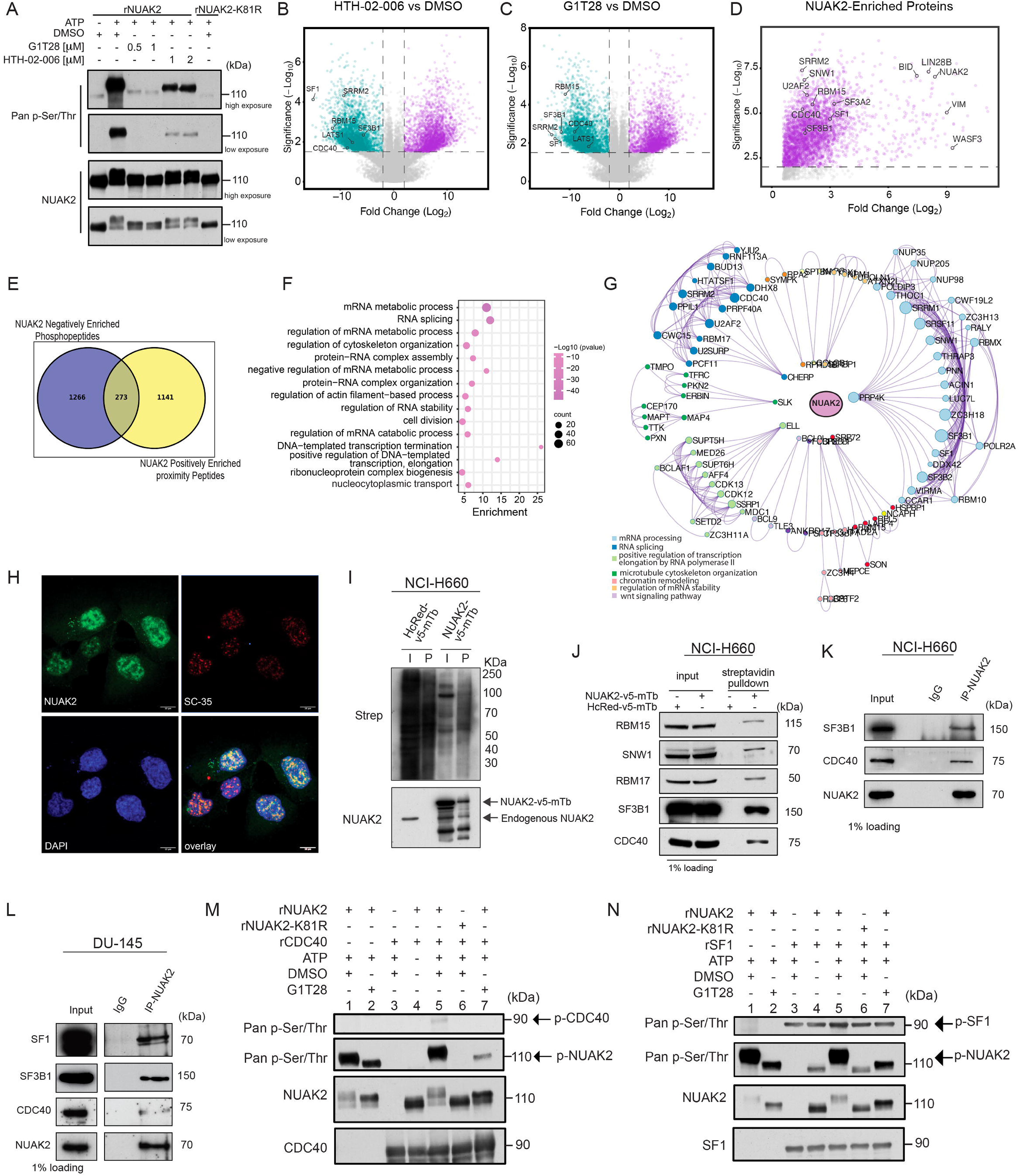
NUAK2 associates with RNA-processing and spliceosome complexes. (A) Representative immunoblot of an in vitro kinase assay assessing rNUAK2 autophosphorylation in the presence of G1T-28 or HTH-02-006, with rNUAK2-K81R (kinase-dead) included as a negative control. NUAK2 autophosphorylation was detected using a pan-phosphoserine/threonine antibody. (B–C) Volcano plots showing differential phosphopeptides in NCI-H660 cells treated with the NUAK2 inhibitors HTH-02-006 (5 µM) or G1T-28 (5 µM) for 1 h compared with DMSO. (D) NUAK2-enriched proteins identified by comparing biotinylated proteins in NUAK2-V5-mTb versus HcRed-V5-mTb (control) samples. (E) Venn diagram depicting overlap of proteins differentially phosphorylated (log₂FC ≤ −5, P ≤ 0.01) and significantly enriched (≥ 2-fold, P ≤ 0.01) relative to control, identifying 273 candidate NUAK2-associated substrate proteins. (F) Enrichment analysis of overlapping proteins highlights mRNA metabolic processes, cytoskeletal organization, mRNA transport, and chromosome organization. (G) Protein–protein interaction network of the 273 overlapping candidate proteins constructed using STRING, BioGRID, OmniPath, and InWeb_IM interaction data, restricted to experimentally validated interactions, analyzed using Molecular Complex Detection (MCODE) to identify densely connected subnetworks, and visualized in Cytoscape. (H) Immunofluorescence staining of NUAK2 and the nuclear speckle marker SC-35 in DU-145 cells demonstrates that NUAK2 is predominantly enriched in nuclear speckles. (I) Validation immunoblots of biotin labeling and affinity pulldown of NUAK2-V5-mTb and HcRed-V5-mTb in NCI-H660 cells. Biotinylated proteins were enriched using streptavidin affinity capture and detected by streptavidin blotting and immunoblotting for NUAK2. (J) Immunoblot validation of NUAK2 proximity interactions in NCI-H660 cells expressing NUAK2-V5-mTb or HcRed-V5-mTb controls. Cells were treated with 0.5 mM biotin for 4 h, followed by streptavidin affinity pulldown to enrich biotinylated proteins. (K) Representative immunoblots of Co-IPs in NCI-H660 cells and (L) DU-145 cells validating the association of endogenous NUAK2 with core splicing factors. (M) Representative immunoblots of in vitro kinase assays evaluating rCDC40 and (N) rSF1 as direct substrates of NUAK2. rCDC40 (M) or rSF1 (N) were incubated with ATP, rNUAK2, rNUAK2-K81R, or G1T-28 as denoted. Substrate phosphorylation was detected using a pan-phosphoserine/threonine antibody, with total protein levels shown as loading controls.

To define NUAK2-associated signaling networks, we employed an integrated proteomic strategy combining phosphoproteomic profiling with complementary biotin-based proximity-labeling proteomics (Fig. S3B). We performed phosphoproteomic profiling of NCI-H660 cells treated with HTH-02-006 or G1T-28. Comparative analysis revealed substantial overlap in differential phosphopeptide abundance induced by the two compounds (Fig. 3B–C, Table S3). Gene ontology analysis of differentially phosphorylated proteins demonstrated significant enrichment of biological processes related to RNA metabolism, RNA processing, and chromatin organization (Fig. S3C–D).

To identify proteins that associate with NUAK2, we performed proximity-labeling proteomics using a NUAK2-miniTurboBioID fusion protein followed by streptavidin affinity purification and mass spectrometry (Fig. S3E–F). This approach identified 1,414 proteins significantly enriched within the NUAK2 proximal interactome relative to control samples (fold change > 2, P < 0.01) (Fig. 3D; Fig. S3G). Functional enrichment analysis of NUAK2-proximal proteins revealed significant enrichment of pathways related to RNA splicing, mRNA processing, and chromatin regulation (Fig. S3H–I).

To prioritize candidate phospho-substrates and NUAK2-associated proteins, we integrated the phosphoproteomic and proximity-labeling datasets, focusing on proteins that exhibited both reduced phosphorylation following NUAK2 inhibitor treatment (log₂FC ≤ −5, P ≤ 0.01) and enrichment in the NUAK2 proximal interactome (log₂FC ≥ 2, P ≤ 0.01). This integrated analysis identified 273 proteins as potential NUAK2 proximal phospho-substrates (Fig. 3E; Table S5). Functional enrichment analysis of these candidates revealed enrichment of pathways related to mRNA metabolism, RNA splicing, cytoskeletal organization, and mRNA transport (Fig. 3F). Notably, more than 100 candidates mapped to RNA-processing pathways, including spliceosome function, transesterification-dependent RNA splicing, and protein–RNA complex organization. Protein–protein interaction network analysis of the overlapping candidates further identified mRNA processing and RNA splicing as prominent functional nodes within the NUAK2-associated network (Fig. 3G). Several key spliceosomal and RNA-processing proteins, including SF1, SF3B1, CDC40, and RBM15, were among candidate NUAK2 proximal phospho-substrates.

Consistent with these findings, immunofluorescence analysis demonstrated predominant nuclear localization of NUAK2 in DU-145 cells, with a punctate staining pattern that overlapped with SC-35, a canonical marker of nuclear speckles that are enriched in splicing factors (Fig. 3H). To validate the association of NUAK2 with spliceosome proteins, we performed streptavidin affinity pulldown following NUAK2-miniTurboBioID proximity labeling and co-immunoprecipitation assays. Streptavidin pulldown followed by immunoblotting confirmed enrichment of multiple spliceosomal and RNA-processing proteins, including SF3B1, CDC40, SNW1, RBM15, and RBM17, within the NUAK2 proximal interactome (Fig. 3I–J). Furthermore, co-immunoprecipitation of endogenous NUAK2 confirmed its association with endogenous SF1, SF3B1, and CDC40 in NCI-H660 and DU-145 cells (Fig. 3K–L).

We next examined whether NUAK2-associated spliceosomal proteins were regulated through NUAK2-dependent phosphorylation. Phosphoproteomic analysis revealed significantly reduced phosphorylation of SF1 at S80 and CDC40 at S43 following treatment with either HTH-02-006 or G1T-28, while both proteins were also highly enriched in the NUAK2 proximity-labeling dataset (Fig. S3J–K). To determine whether SF1 and CDC40 are direct substrates of NUAK2, we performed in vitro kinase assays using rNUAK2 and rNUAK2-KD. Both proteins were robustly phosphorylated in the presence of rNUAK2, whereas phosphorylation was markedly reduced by G1T-28 treatment and was absent in reactions containing rNUAK2-KD (Fig. 3M–N). These findings provide qualitative evidence that SF1 and CDC40 are direct NUAK2 phospho-substrates.

Collectively, our integrated phosphoproteomic, proximity-labeling, interaction, and biochemical analyses identify NUAK2 as a kinase associated with spliceosomal and RNA-processing machinery and support a mechanistic role for NUAK2-dependent phosphorylation in regulating RNA-processing programs.

### NUAK2 perturbation disrupts oncogenic RNA-splicing programs in aggressive PCa

To determine whether NUAK2-associated spliceosome interactions translate into functional changes in RNA processing, we next performed transcriptomic profiling following pharmacologic perturbation using G1T-28. RNA sequencing of NCI-H660 cells treated with G1T-28 revealed widespread alterations in gene expression, with 3,611 genes upregulated and 1,907 genes downregulated (|log₂FC| ≥ 2, P ≤ 0.01) (Fig. 4A, Table S6). Principal component analysis confirmed robust transcriptional separation between control and treated cells (Fig. S4A). Gene set enrichment analysis (GSEA) of differentially expressed genes revealed significant enrichment of pathways associated with RNA processing, mRNA splicing, and chromatin regulation (Fig. S4B–C). Notably, spliceosome-related gene sets were among the most significantly enriched pathways.

**Figure 4.**
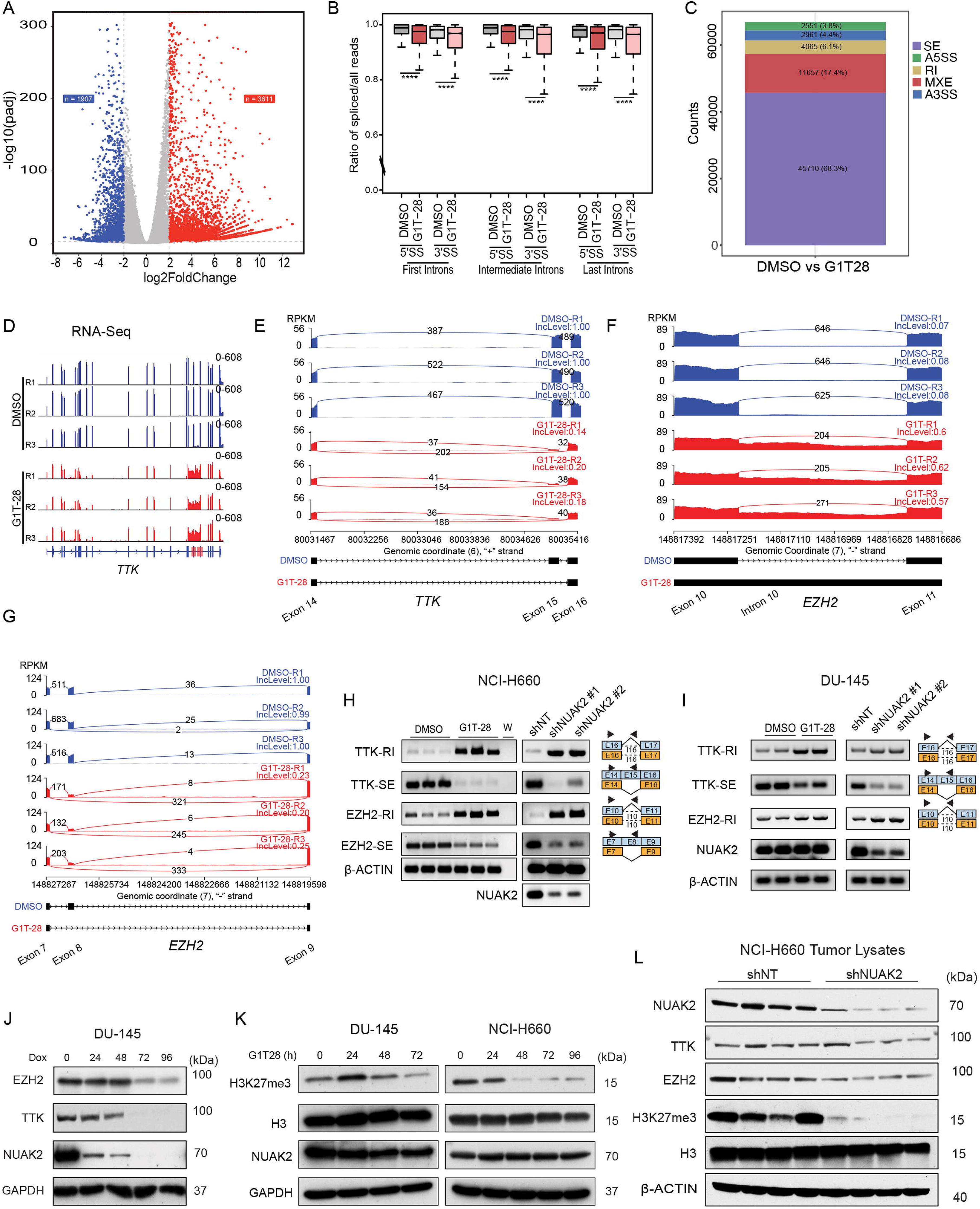
NUAK2 perturbation disrupts RNA splicing of oncogenic regulators. (A) Volcano plot of differentially expressed genes in NCI-H660 cells treated with G1T-28 (5 µM) or DMSO for 24 h. (B) Box plots of spliced-to-total read ratios stratified by intron position (first, middle, last) in RNA-seq of NCI-H660 cells treated with DMSO or G1T-28 for 24 h, across 8,518 isoforms. Statistical significance was assessed by Wilcoxon signed-rank test (**** P < 2.2 × 10⁻¹). (C) Number of differential splicing events between DMSO- and G1T-28-treated NCI-H660 cells, depicting the types and counts identified by rMATS. (D) IGV tracks showing RNA-seq read coverage across the *TTK* locus in NCI-H660 cells treated with G1T-28 or DMSO (control) for 24 h. (E–G) Sashimi plots depicting splicing alterations in *TTK* (SE) and *EZH2* (RI and SE) following G1T-28 treatment. (H) RT–PCR and agarose gel electrophoresis for validation of *TTK* and *EZH2* splicing alterations in NCI-H660 cells following treatment with G1T-28 (5 µM, 24 h) or doxycycline-induced NUAK2 knockdown for 72 h. (I) RT–PCR and agarose gel electrophoresis for validation of *TTK* and *EZH2* splicing alterations in DU-145 cells following G1T-28 treatment (2 µM, 6 h) or doxycycline-induced NUAK2 knockdown (60 h). (J) Immunoblot analysis of EZH2 and TTK in DU-145 cells following time-course doxycycline-induced NUAK2 knockdown. (K) Immunoblot analysis of DU-145 and NCI-H660 cells treated with G1T-28 (1 µM) for the indicated time points. (L) Immunoblot analysis of NCI-H660 CDX tumor lysates expressing shNT or shNUAK2, demonstrating altered EZH2, TTK, and H3K27me3 levels following NUAK2 depletion, with NUAK2 immunoblotting confirming target knockdown.

Given the identification and validation of NUAK2 interactions with RNA-splicing proteins in mass spectrometry-based studies, we analyzed RNA-seq data for global splicing alterations upon NUAK2 perturbation, as described previously (48). Calculation of splicing ratios for 5′ and 3′ splice sites revealed a marked accumulation of unspliced intronic reads across the transcriptome upon NUAK2 inhibition, particularly when introns were stratified by position within transcripts (first, middle, or last) (Wilcoxon signed-rank test, P < 2.2 × 10⁻¹), indicating NUAK2 inhibition globally impairs pre-mRNA splicing efficiency (Fig. 4B and Fig. S4D).

We next performed rMATS analysis of alternative splicing, which identified widespread splicing alterations following NUAK2 perturbation across multiple event categories, including skipped exons (SE), alternative 5′ splice sites (A5SS), alternative 3′ splice sites (A3SS), mutually exclusive exons (MXE), and retained introns (RI) (Fig. 4C). Exon-skipping events represented the most abundant class of alternative splicing alterations (Fig. S4E). Gene Ontology (GO) analysis of alternatively spliced genes revealed significant enrichment of pathways related to the mitotic cell cycle, chromatin regulation, and RNA metabolism (Fig. S4F). Consistently, KEGG pathway analysis showed enrichment of cell cycle–related pathways, Polycomb repressive complex–associated signaling, and pathways linked to neurodegenerative diseases (Fig. S4G).

We next examined genes exhibiting NUAK2-associated splicing alterations within GO and KEGG pathways related to cell-cycle progression, mitosis, and chromatin regulation, processes implicated in tumor growth and lineage plasticity (54, 55). Among these functionally enriched pathways, we prioritized *TTK* and *EZH2* for further validation. *TTK*, encoding a mitotic checkpoint kinase, was prominently represented within mitotic cell-cycle and spindle-organization pathways, whereas *EZH2*, encoding the catalytic histone methyltransferase of PRC2, was enriched within pathways related to chromatin organization and Polycomb repressive complex signaling. Both genes have established roles in cancer progression and therapeutic resistance, with EZH2 particularly implicated in NEPC development (56, 57).

We next performed correlation analyses between *NUAK2* transcript expression and *TTK* or *EZH2* expression in independent PCa cohorts. In the Beltran and Labrecque cohorts, *NUAK2* expression correlated with *TTK* (r = 0.709 and 0.63, P < 0.05) and *EZH2* (r = 0.762 and 0.66, P < 0.05) (Fig. S4H–K) (9, 29). Moreover, EZH2 activity scores positively correlated with *NUAK2* expression in both cohorts (r = 0.684 and 0.593; P < 0.05), further linking *NUAK2* expression with EZH2-associated activity in PCa (Fig. S4L–M) (57).

Manual inspection of *TTK* and *EZH2* transcripts in the Integrative Genomics Viewer (IGV) corroborated splicing alterations following G1T-28 treatment, including intron-retention and exon-skipping events (Fig. 4D, Fig. S4N). Consistently, rMATS sashimi plots demonstrated distinct changes in splice-junction usage within *TTK* and *EZH2* transcripts following G1T-28 treatment (Fig. 4E–G). We next validated these splicing alterations by RT-PCR in both NCI-H660 and DU-145 cells, using primers positioned as indicated across retained-intron or skipped-exon regions (Fig. 4H–I). To determine whether these events were specifically associated with NUAK2, we performed parallel genetic depletion of NUAK2 using doxycycline-inducible shRNAs. NUAK2 knockdown recapitulated the *TTK* and *EZH2* splicing alterations observed following G1T-28 treatment, providing orthogonal genetic evidence that NUAK2 regulates the splicing of these transcripts (Fig. 4H–I).

We next examined the functional consequences of NUAK2 perturbation on TTK and EZH2 protein expression. NUAK2 depletion in DU-145 cells, as well as G1T-28 treatment in DU-145, NCI-H660, and PARCB-1 cells, resulted in progressive reductions in TTK and EZH2 protein levels (Fig. 4J; Fig. S4O). G1T-28 treatment also reduced H3K27me3 levels, consistent with diminished EZH2 activity (Fig. 4K). Importantly, these effects were recapitulated in vivo, where NUAK2-depleted NCI-H660 tumors exhibited reduced TTK and EZH2 protein expression together with decreased H3K27me3 levels (Fig. 4L). Collectively, these findings demonstrate that NUAK2 regulates *TTK* and *EZH2* RNA processing and links NUAK2-dependent splicing alterations to downstream changes in protein expression and EZH2-associated chromatin regulation.

### NUAK2 is pharmacologically tractable and engaged by clinically relevant inhibitors

To evaluate the therapeutic potential of NUAK2 inhibition, we first examined the activity of the semi-selective preclinical NUAK2 inhibitor HTH-02-006 *in vitro* (53). HTH-02-006 induced a dose-dependent growth inhibition in NCI-H660, PARCB-1, and DU-145 cells, with IC₅₀ values in the low micromolar range (Fig. 5A). Consistently, HTH-02-006 significantly suppressed DU-145-RFP spheroid growth (Fig. 5B–C) and reduced clonogenic potential in a dose-dependent manner (Fig. 5D–E).

**Figure 5.**
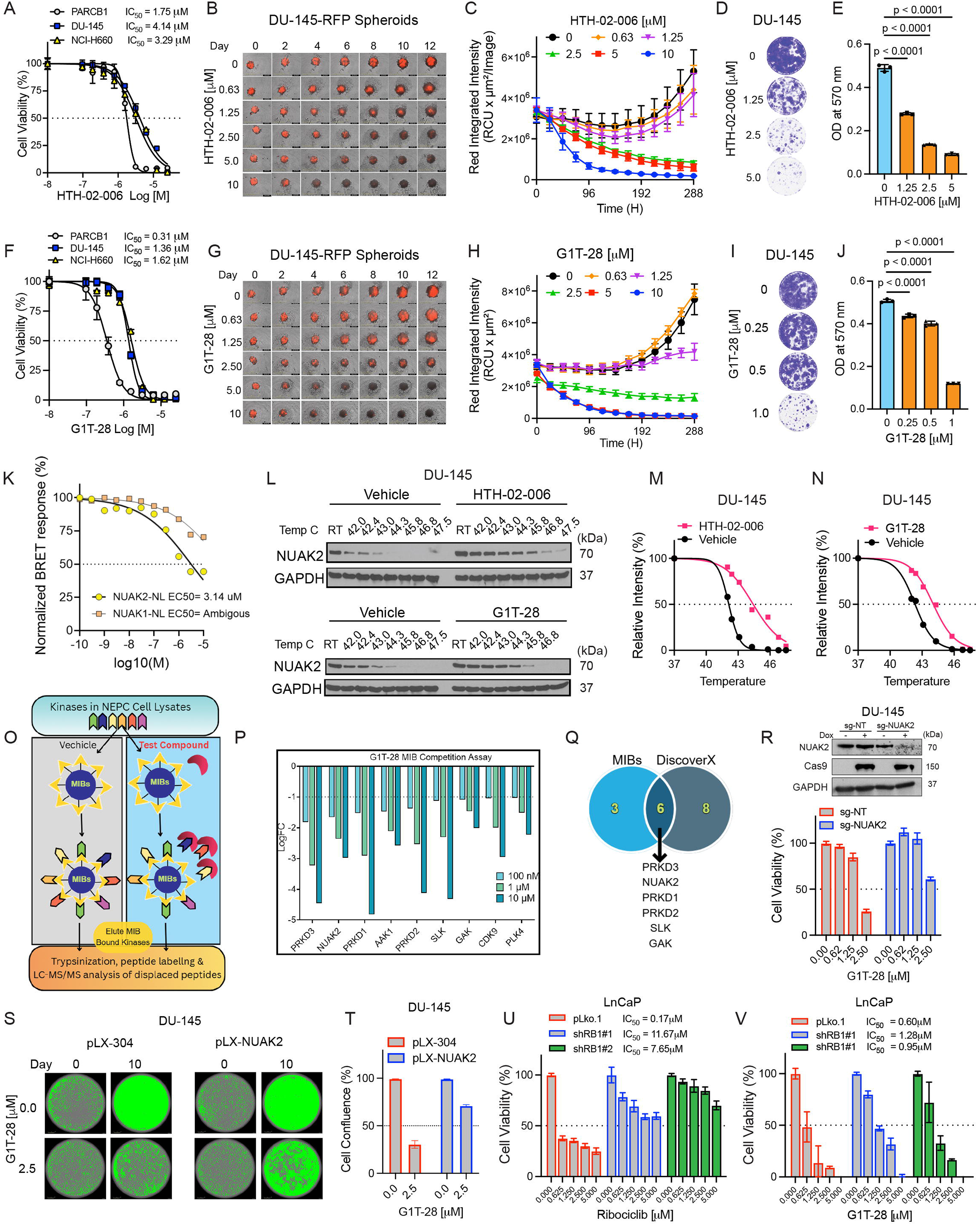
Clinically relevant inhibitors engage NUAK2 and suppress AR-indifferent prostate cancer cell growth. (A) Dose response curves of NCI-H660, PARCB-1, and DU-145 upon treatment with HTH-02-006. Data represent mean ± SD from three independent biological replicates. (B) Representative images and (C) quantification of DU-145-RFP spheroid growth kinetics following HTH-02-006 treatment. Data are shown as mean ± SEM from three biological replicates. (D) Representative images and (E) quantification of DU-145 colony formation assay after HTH-02-006 treatment. Data represent mean ± SD from three biological replicates. (F) Dose response of NCI-H660, PARCB-1, and DU-145 following treatment with G1T-28. Data represent mean ± SD from three biological replicates. (G) Representative images and (H) quantification of growth kinetics of DU-145-RFP spheroids treated with G1T-28. Data represent mean ± SEM from three biological replicates. (I) Representative images and (J) quantification of DU-145 colony formation assay after G1T-28 treatment. Data represent mean ± SD from three biological replicates. (K) G1T-28-NanoBRET intracellular kinase assay using NUAK2-NanoLuc and NUAK1-NanoLuc fusion proteins in 293T cells. (L) Immunoblot analysis of cellular thermal shift assay (CETSA) assessing NUAK2 stability in DU-145 treated with DMSO (vehicle), HTH-02-006 (25 µM), or G1T-28 (6.25 µM). (M–N) NUAK2 thermal stability and melting curves generated from CETSA immunoblot analyses of DU-145 cells following treatment with HTH-02-006 or G1T-28. (O) Schematic overview of the MIB competition assay workflow performed with PARCB-1 cell lysates followed by mass spectrometry analysis. (P) Kinases displaced from MIBs by G1T-28 with log₂FC ≤ −1 at 100 nM, 1 μM, and 10 μM. (Q) Venn diagram illustrating the overlap between kinases identified in the reported G1T-28 DiscoveRx kinome scan and those captured by the MIB assay. (R) Western blot and cell viability analysis of TRE3G-Cas9-DU-145 cells expressing sgNT or sgNUAK2 following treatment with G1T-28 at the indicated concentrations for 96 h. (S) Representative images and (T) quantification of cell confluence analysis of DU-145 transduced with empty vector pLX-304 control or pLX-NUAK2 treated with G1T-28 as indicated. (U) Cell viability of LNCaP cells expressing pLKO.1 control or independent RB1-targeting shRNAs (shRB1 #1 and shRB1 #2) treated with ribociclib as indicated for 96 h. (V) Cell viability of LNCaP cells expressing pLKO.1 control or independent RB1-targeting shRNAs (shRB1 #1 and shRB1 #2) treated with G1T-28 as indicated for 96 h. Data represent mean ± SD from three independent biological replicates. IC_50_ and melting temperature were calculated using non-linear regression (curve fit) for panels (A, F, K, M, and N). Statistical analyses were performed using two-way ANOVA for panels (C and H), and one-way ANOVA with Tukey’s multiple-comparison test for (E and J).

As described above, interrogation of the Pharos database identified the CDK4/6 inhibitors G1T-28 (trilaciclib) and G1T-38 (lerociclib) among compounds with reported high-affinity binding to NUAK2 (Fig. S3A). Previously published KINOMEscan profiling demonstrated >90% inhibition of NUAK2 at 100 nM by both compounds (Fig. S5A–B) (58, 59). Thus, we next evaluated the antitumor activity of G1T-28 and G1T-38 in PCa cells. We prioritized G1T-28 for subsequent studies because of its FDA approval, while G1T-38 remains under clinical development and has recently received regulatory approval for the treatment of breast cancer in China (60).

*In vitro*, G1T-28 and G1T-38 induced robust, dose-dependent growth inhibition in NCI-H660, PARCB-1, and DU-145 cells despite *RB1* loss or mutation, suggesting activity independent of canonical CDK4/6–RB signaling (Fig. 5F; Fig. S5C). Consistently, both compounds markedly suppressed DU-145 spheroid growth in 3D culture (Fig. 5G–H; Fig. S5D–E) and significantly reduced clonogenic capacity (Fig. 5I–J; Fig. S5F–G). In contrast, ribociclib, another FDA-approved CDK4/6 inhibitor with a different chemical structure and minimal NUAK2 activity, exhibited limited activity in NCI-H660, PARCB-1, and DU-145 cells at comparable drug concentrations, consistent with their RB-deficient status and lack of ribociclib-NUAK2 engagement (61) (Fig. S5H).

To further validate G1T-28 engagement of NUAK2 in cells, we performed an intracellular NanoBRET target-engagement assay using a NUAK2–NanoLuc fusion construct. G1T-28 displaced the NanoBRET tracer in a dose-dependent manner, with an estimated IC₅₀ of 3.14 μM, demonstrating intracellular engagement of NUAK2 (Fig. 5K). In contrast, no measurable engagement of NUAK1 was observed. CETSA confirmed engagement of endogenous NUAK2 by HTH-02-006, as evidenced by pronounced thermal stabilization of NUAK2 in DU-145 cells (Fig. 5L–N). Importantly, G1T-28 similarly stabilized NUAK2, further supporting direct cellular target engagement (Fig. 5L–N). Consistent effects were observed in 22Rv1 cells, where both G1T-28 and G1T-38 induced thermal stabilization of NUAK2 (Fig. S5I–K). Furthermore, in RB-positive LNCaP cells, G1T-28 stabilized both CDK4 and NUAK2, demonstrating concurrent cellular engagement of its canonical CDK4/6 target and NUAK2 within the same cellular context (Fig. S5L–N).

To further characterize the kinase-engagement profile of G1T-28 in an unbiased manner, we performed multiplexed inhibitor bead (MIB) competition profiling coupled with mass spectrometry (Fig. 5O). In vehicle-control samples, MIBs captured 412 kinases from PARCB-1 cell lysates (Table S2). G1T-28 induced dose-dependent displacement of NUAK2, with log₂ fold changes of −1.65, −2.36, and −2.98 at 100 nM, 1 μM, and 10 μM, respectively, further supporting NUAK2 engagement (Fig. 5P). Consistent with the NanoBRET results, NUAK1 was not displaced. G1T-28 exhibited a relatively restricted kinase-engagement profile at 100 nM, with only three kinases displaying log₂FC ≤ −1.5. Although CDK4/6 were captured by MIBs in control samples, neither kinase was detectably displaced by G1T-28 in PARCB-1 lysates. At higher concentrations, G1T-28 also engaged several previously reported kinase targets, including PRKD1/2/3, GAK, and SLK (Fig. 5Q).

To test the relative contribution of NUAK2 inhibition to the growth-suppressive activity of G1T-28, we next examined cellular response to G1T-28 following genetic perturbation of NUAK2. Because complete NUAK2 depletion was lethal in these models, precluding subsequent drug-response analysis, we generated cells with partial NUAK2 loss. Partial NUAK2 knockout reduced sensitivity to G1T-28, consistent with reduced dependence on the available drug target (Fig. 5R; Fig. S5O). Conversely, NUAK2 overexpression in DU-145 cells imparted G1T-28 resistance (Fig. 5S–T), supporting NUAK2 as a functionally relevant target contributing to G1T-28 activity. To further distinguish NUAK2-dependent activity from canonical CDK4/6 inhibition, we compared G1T-28 and G1T-38 with ribociclib in RB1-positive LNCaP cells following RB1 shRNA knockdown. As expected, RB1 depletion rendered LNCaP cells resistant to ribociclib, consistent with the requirement for an intact RB pathway for canonical CDK4/6 inhibitor activity (61) (Fig. 5U). In contrast, RB1-depleted LNCaP cells, with intact NUAK2 expression, remained sensitive to both G1T-28 and G1T-38 (Fig. 5V; Fig. S5P).

Collectively, our biochemical and genetic studies demonstrate that these clinically advanced compounds directly engage NUAK2 and that NUAK2 inhibition contributes to the antitumor activity of G1T-28 and G1T-38 in RB-deficient prostate cancer. These findings establish NUAK2 as a pharmacologically tractable therapeutic target.

### NUAK2 inhibition suppresses tumor growth and enhances chemotherapy response

To determine whether NUAK2 inhibition confers therapeutic benefit *in vivo*, we evaluated both the preclinical NUAK2 inhibitor HTH-02-006 and the clinically approved compound G1T-28 in multiple NEPC models (Fig. S6A). In cell line–derived xenograft (CDX) studies, HTH-02-006 significantly suppressed tumor growth kinetics and endpoint tumor weights in NCI-H660 (Fig. 6A–B) and PARCB-1 models (Fig. 6C–D). Consistent effects were observed in the LTL-352 NEPC PDX model, where HTH-02-006 markedly inhibited tumor growth kinetics and endpoint tumor weights (Fig. 6E–F). HTH-02-006 treatment was well tolerated, with no significant changes in body weight throughout the treatment period (Fig. S6B–D).

**Figure 6.**
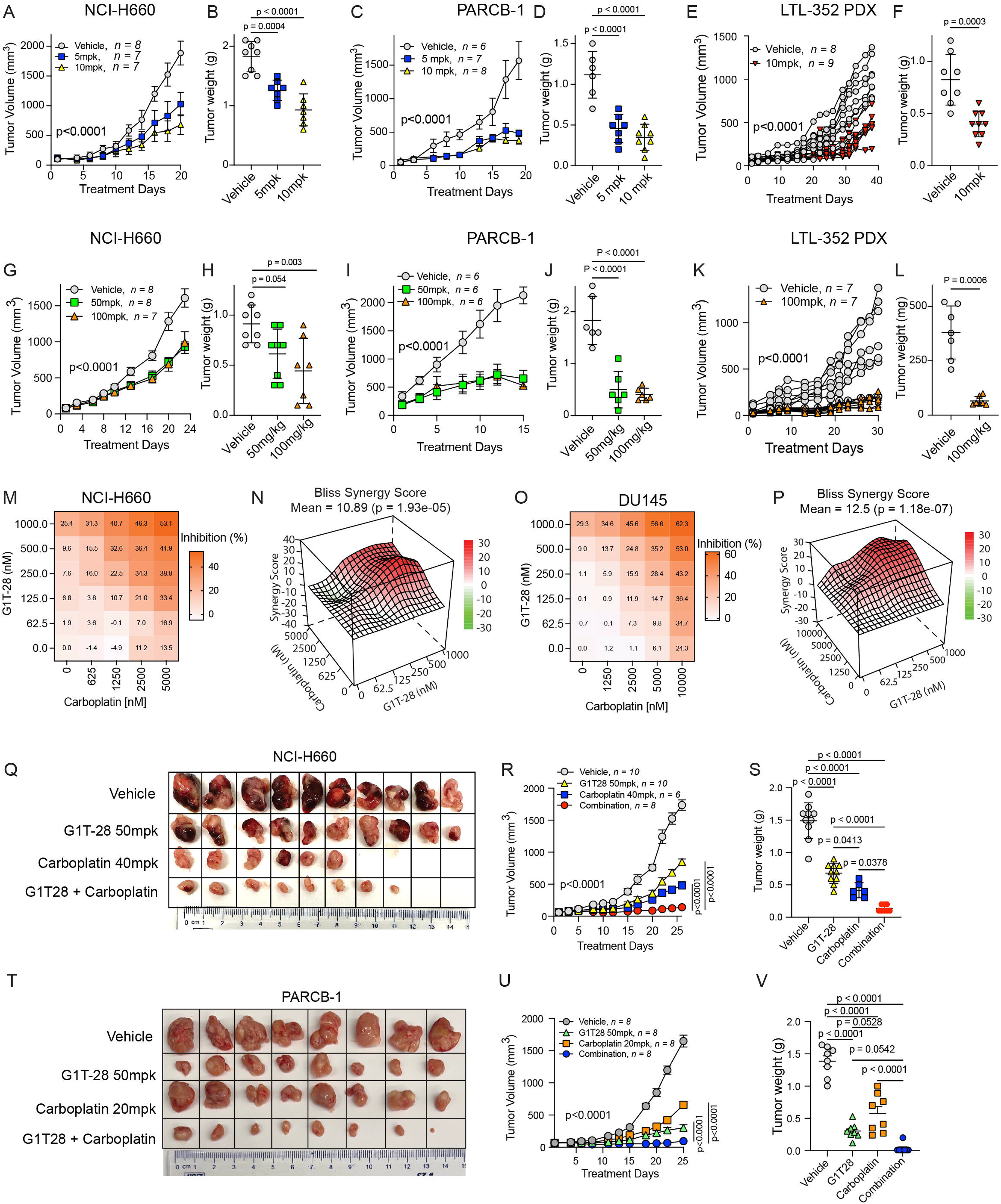
Pharmacologic targeting of NUAK2 suppresses tumor growth and potentiates chemotherapy response. (A) NCI-H660 CDX longitudinal tumor growth kinetics and (B) endpoint tumor weights from mice treated with vehicle (n = 8), HTH-02-006 at 5 mpk (n = 7), or HTH-02-006 at 10 mpk (n = 7). (C) PARCB-1 CDX longitudinal tumor growth kinetics and (D) endpoint tumor weights from mice treated with vehicle (n = 6), HTH-02-006 at 5 mpk (n = 7), or HTH-02-006 at 10 mpk (n = 8). (E) LTL-352 NEPC PDX tumor growth kinetics and (F) endpoint tumor weights from mice treated with vehicle (n = 8) or HTH-02-006 (10 mpk; n = 9). (G) NCI-H660 CDX longitudinal tumor volume measurements and (H) endpoint tumor weights from mice treated with vehicle (n = 8), G1T-28 at 50 mpk (n = 9), or G1T-28 at 100 mpk (n = 9) at treatment initiation. At endpoint, n = 8, 8, and 7 mice remained in the vehicle, 50 mpk, and 100 mpk groups, respectively. (I) PARCB-1 CDX longitudinal tumor volume measurements and (J) endpoint tumor weights from mice treated with G1T-28 (50 or 100mpk) or vehicle (n = 6 mice per group). (K) LTL-352 NEPC PDX longitudinal tumor volume measurements and (L) endpoint tumor weights from mice treated with vehicle (n = 8) or G1T-28 (100 mpk; n = 10) at treatment initiation. At endpoint, n = 7 mice remained in each group. (M) NCI-H660 G1T-28 and carboplatin combination matrix growth inhibition heatmap and (N) Bliss synergy plots. (O) DU-145 G1T-28 and carboplatin combination matrix growth inhibition heatmap and (P) Bliss synergy plots. (Q) Representative images of excised NCI-H660 CDX tumors, (R) tumor growth kinetics, and (S) endpoint tumor weights from mice treated with vehicle, G1T-28 (50 mpk), carboplatin (40 mpk), or the combination. Mice were randomized at n = 10 per group at treatment initiation. At endpoint, n = 6 mice remained in the carboplatin group and n = 8 in the combination group. (T) Representative images of excised PARCB-1 CDX tumors, (U) tumor growth kinetics, and (V) endpoint tumor weights from mice treated with vehicle, G1T-28 (50 mpk), carboplatin (20 mpk), or the combination (n = 8 mice per group). mpk – mg/kg; statistical analyses were performed using two-way ANOVA for panels A, C, E, G, I, K, R and U, and one-way ANOVA for panels B, D, F, H, J, L, S and V.

We next evaluated the therapeutic efficacy of G1T-28 *in vivo*. In NCI-H660 xenografts, G1T-28 significantly suppressed tumor growth kinetics and endpoint tumor weights (Fig. 6G–H), with mean tumor volumes of 1,607.7 mm³ in vehicle-treated mice compared with 931.4 mm³ and 990.6 mm³ in mice treated with 50 and 100 mg/kg G1T-28, respectively. PARCB-1 xenografts exhibited greater sensitivity, with >75% inhibition of tumor growth and a corresponding reduction in tumor weight, relative to vehicle-treated controls (Fig. 6I–J). G1T-28 also demonstrated robust efficacy in the LTL-352 NEPC PDX model, resulting in near-complete suppression of tumor growth (Fig. 6K–L). G1T-28 was well tolerated across all three models, with no significant changes in body weight during treatment (Fig. S6E–G).

Given the activity of G1T-28 as a monotherapy, we next investigated whether NUAK2 inhibition could enhance the efficacy of standard chemotherapy used clinically for patients with NEPC. In 5 × 6 dose-matrix assays, G1T-28 synergized with carboplatin in both NCI-H660 and DU-145 cells, with mean Bliss synergy scores of 10.89 and 12.5, respectively (Fig. 6M–P). Synergistic activity was also observed with docetaxel in both cell lines (Fig. S6H–K). We subsequently evaluated the combination of G1T-28 plus carboplatin in vivo using NCI-H660 xenografts treated with vehicle, G1T-28 (50 mg/kg, once daily), carboplatin (40 mg/kg, once weekly), or the combination. G1T-28 and carboplatin monotherapies reduced tumor growth by 51.3% and 72.3%, respectively, whereas the combination achieved 91.9% tumor growth inhibition and significantly reduced endpoint tumor weights relative to either monotherapy (Fig. 6Q–S). Both single-agent and combination treatments were well tolerated, with no significant changes in body weight during treatment (Fig. S6L). We further performed complete blood count (CBC) analysis to assess potential hematologic toxicity and evaluate the myeloprotective effects of G1T-28. However, carboplatin did not induce neutropenia in mice, consistent with previous reports (62, 63), precluding a direct assessment of the myeloprotective effects of G1T-28 in this setting (Fig. S6M). Furthermore, consistent results were observed in the PARCB-1 xenograft model, where the combination treatment resulted in enhanced tumor growth inhibition relative to single-agent therapy, despite a reduction in the carboplatin dose to 20 mg/kg (Fig. 6T–V). Notably, PARCB-1 tumors displayed greater sensitivity to G1T-28 than NCI-H660 xenografts, mirroring earlier *in vivo* results (Fig. 6I–J).

Collectively, these results demonstrate that NUAK2 inhibition suppresses tumor growth and enhances chemotherapy response across multiple preclinical models of AR-indifferent PCa.

## Discussion

The widespread use of second- and third-generation AR-targeted therapies has contributed to the emergence of AR-indifferent mCRPC, including subtypes such as DNPC and NEPC. These subtypes represent highly lethal disease states characterized by rapid progression and limited treatment options. Patients with treatment-emergent NEPC and DNPC present with extensive metastatic disease and have a median survival of less than one year (7, 64). Platinum-based chemotherapy, often combined with etoposide, remains a mainstay of treatment for NEPC; however, responses are typically short-lived and often accompanied by substantial toxicity, including myelosuppression and impaired quality of life (18). Thus, there remains a critical need to identify therapeutically tractable molecular targets for AR-indifferent PCa.

Our findings nominate NUAK2 as a previously unrecognized target in NEPC and a subset of DNPC tumors. Genetic depletion of NUAK2 suppressed proliferation, clonogenic growth, and tumor progression across multiple AR-indifferent PCa models, whereas NUAK2 overexpression enhanced tumor cell growth. Importantly, these effects were predominantly dependent on NUAK2 kinase activity. Although the kinase-dead NUAK2 mutant moderately enhanced PC-3 proliferation in vitro, it failed to enhance clonogenic growth and exhibited markedly reduced metastatic potential in orthotopic models, indicating that NUAK2 kinase activity is particularly important for sustained tumor progression. Nevertheless, the residual activity observed in certain in vitro contexts raises the possibility of kinase-independent functions that warrant further investigation.

Although NUAK2 has been implicated in several cancer types, its molecular functions remain poorly defined, and few direct substrates or interacting partners have been characterized (65). Our integrative proteomic analysis provided a framework for defining NUAK2-regulated signaling networks and revealed an unexpected connection between NUAK2 and RNA-processing machinery. More than 250 proteins were both proximal to NUAK2 and differentially phosphorylated following pharmacologic NUAK2 perturbation, with enrichment of spliceosomal and RNA-processing proteins, including SF3B1, CDC40, and SF1. Consistent with these findings, NUAK2 localized predominantly to the nucleus and associated with multiple components of the spliceosomal machinery. Moreover, phosphoproteomic analysis identified reduced phosphorylation of SF1 and CDC40 following NUAK2 inhibition, while biochemical kinase assays demonstrated phosphorylation of both proteins by rNUAK2. Importantly, substrate phosphorylation was markedly reduced by G1T-28 and absent with rNUAK2-KD, providing additional evidence linking NUAK2 kinase activity to spliceosomal proteins. Together, these findings identify a previously unrecognized association between NUAK2 signaling and RNA-processing machinery.

Consistent with this mechanistic connection, pharmacologic NUAK2 perturbation resulted in widespread alterations in RNA-splicing programs, including changes in genes involved in cell-cycle regulation, mitosis, and chromatin organization. Dysregulated RNA processing has emerged as a critical driver of tumor progression, lineage plasticity, and therapeutic resistance in cancer, including PCa. Pharmacologic targeting of spliceosome components is therefore being actively explored, with several RNA-splicing inhibitors currently in clinical trials for cancer treatment (66, 67). Malignancies harboring mutations in splicing factors such as SF3B1, SRSF2, or U2AF1 have demonstrated vulnerabilities to perturbation of spliceosome function (68, 69). Our findings suggest that NUAK2 may provide an alternative approach for therapeutically perturbing RNA splicing in tumors lacking canonical spliceosome mutations. The NUAK2-dependent splicing alterations observed in TTK and EZH2, together with corresponding changes in protein expression and H3K27me3, further connect NUAK2 signaling with mitotic and epigenetic programs implicated in aggressive PCa. Thus, NUAK2 may function as a regulatory node linking kinase signaling to RNA-processing networks that support tumor progression.

Beyond its mechanistic role, our findings demonstrate that NUAK2 is pharmacologically tractable. Biochemical kinase assays showed that G1T-28 directly inhibited NUAK2 autophosphorylation and NUAK2-mediated phosphorylation of CDC40 and SF1. Complementary NanoBRET and CETSA assays demonstrated cellular engagement of NUAK2, while kinome-wide MIB competition profiling in NEPC cell lysates further supported NUAK2 engagement and characterized the broader kinase-binding profile of G1T-28. Importantly, genetic perturbation provided additional functional evidence linking NUAK2 to G1T-28 activity. NUAK2 loss reduced sensitivity to G1T-28, whereas NUAK2 overexpression shifted cells toward resistance. Together, these orthogonal biochemical, target-engagement, kinome-profiling, and genetic approaches support NUAK2 as a functionally relevant target of G1T-28.

The activity of G1T-28 and G1T-38 in *RB1*-deficient models further distinguishes their antitumor effects from canonical CDK4/6 inhibition (61, 70). G1T-28 and G1T-38 suppressed growth of *RB1*-deficient or mutant models that were comparatively resistant to ribociclib (61, 70). Furthermore, *RB1* depletion rendered LNCaP cells resistant to ribociclib, whereas sensitivity to G1T-28 and G1T-38 was retained. These findings support an antitumor mechanism for these compounds that extends beyond canonical CDK4/6–RB signaling and is consistent with a contribution from NUAK2 inhibition. The translational potential of these observations is supported by the clinical development of both compounds. G1T-28 is FDA-approved for reducing chemotherapy-induced myelosuppression in extensive-stage SCLC, while G1T-38 remains under clinical development and has recently received regulatory approval for the treatment of breast cancer in China (71, 72). Thus, the identification of NUAK2-directed activity among clinically advanced compounds provides an opportunity for therapeutic repurposing while more selective NUAK2 inhibitors are developed. Nevertheless, because G1T-28 engages additional kinases, the contribution of other targets to its antitumor activity cannot be excluded. Development of more selective NUAK2 inhibitors will therefore be important for further defining the therapeutic window and target-specific effects of NUAK2 inhibition.

Pharmacologic NUAK2 targeting demonstrated substantial antitumor activity *in vivo*. Both HTH-02-006 and G1T-28 suppressed tumor growth across CDX and PDX NEPC models, supporting the therapeutic potential of pharmacologic NUAK2 inhibition. Moreover, G1T-28 enhanced the efficacy of platinum-based chemotherapy, with synergistic activity observed *in vitro* and significantly enhanced tumor growth inhibition with G1T-28 plus carboplatin as compared with either monotherapy *in vivo*. Clinically, G1T-28 is FDA-approved for chemotherapy-induced myelosuppression in extensive-stage small-cell lung cancer (SCLC) and is administered prior to chemotherapy, which is typically given every three weeks (71). Because sustained NUAK2 inhibition may require a different dosing paradigm involving daily treatment, there is a risk of adverse effects. Thus, we evaluated the hematologic effects of daily G1T-28 administration, alone and in combination with carboplatin, following 26 days of treatment. Daily G1T-28 did not adversely affect neutrophil, red blood cell, monocyte, or platelet counts in our models (Fig. S6M). Although carboplatin is well documented to induce neutropenia and thrombocytopenia in patients, it did not reduce neutrophil counts in our preclinical model and instead produced a modest increase, consistent with previous observations (62, 63). Carboplatin did, however, significantly reduce platelet counts, an effect that was slightly attenuated by G1T-28 co-treatment (Fig. S6M). Given the biological and therapeutic similarities between NEPC and SCLC, these findings raise the possibility that G1T-28 could provide complementary therapeutic effects in NEPC by retaining its established myeloprotective properties while exerting tumor-directed activity that includes NUAK2 inhibition.

Several important questions remain. Although our biochemical studies provide qualitative evidence that NUAK2 phosphorylates CDC40 and SF1, the precise NUAK2-dependent phosphorylation sites and their functional consequences for spliceosome activity remain to be rigorously validated. While phosphoproteomic analysis identified reduced phosphorylation of SF1 S80 and CDC40 S43 following pharmacologic perturbation, direct assessment of these specific residues as NUAK2 phosphorylation sites will require site-resolved biochemical validation. The lack of phospho-specific antibodies further limited our ability to characterize these phosphorylation events in cellular models. In addition, whether NUAK2 directly contributes to lineage plasticity and adenocarcinoma-to-neuroendocrine transdifferentiation through regulation of RNA-processing programs remains unresolved. Finally, because G1T-28 engages multiple kinases, development of selective NUAK2 inhibitors and further genetic and pharmacological studies will be necessary to define the extent to which NUAK2 inhibition accounts for its antitumor activity. Addressing these questions will further clarify how NUAK2 integrates kinase signaling with RNA-processing pathways and determine the broader therapeutic potential of targeting this axis in aggressive PCa.

In conclusion, our study identifies NUAK2 as a therapeutic target in AR-indifferent PCa and uncovers a previously unrecognized connection between NUAK2 kinase activity and RNA-processing regulation. Our findings establish the therapeutic tractability of NUAK2, provide evidence that NUAK2 contributes to the antitumor activity of G1T-28 in RB-deficient PCa, and support the development of selective NUAK2-directed therapeutic strategies for aggressive, treatment-refractory disease.

## Supporting information

Table S1. List of primers.

Table S3. Differential phosphorylation analysis following NUAK2 inhibition with G1T-28 and HTH-02-006.

Table S4. Differentially enriched proximity-labeled proteins in NUAK2-V5-miniTurbo versus HcRed-V5-miniTurbo samples.

Table S5. Overlapping candidate NUAK2-associated proteins identified by integrated phosphoproteomic and proximity labeling analysis.

Table S6. Differential gene expression in NCI-H660 cells following G1T-28 treatment.

Table S7. Splicing alterations in NCI-H660 cells upon NUAK2 inhibition identified by rMATS.

Table S8. Correlation of NUAK2 expression with neuroendocrine and prostate adenocarcinoma markers.

## Declarations

### Ethics approval and consent to participate

All animal experiments were performed in accordance with ethical regulations and approved protocols from the Institutional Animal Care and Use Committee (IACUC) of Duke University (Protocol: A190-23-09 and A034-25-04) and the Department of Defense Animal Care and Use Review Office (ACURO).

### Consent for publication

Not applicable.

## Availability of data and materials

The datasets used and/or analyzed during the current study are available from the corresponding author on reasonable request.

## Competing interests

AJA receives research support (to Duke) from the NIH/NCI, PCF, DoD, Astellas, Pfizer, Bayer, Janssen, BMS, AstraZeneca, Merck, Pathos, Amgen, Novartis. AJA declares consulting or advising relationships with Astellas, Pfizer, Bayer, Janssen, BMS, AstraZeneca, Merck, Exelixis, Novartis, Medscape, Telix, Duality Bio, MedIQ, IDEOlogy, Sumitomo, and Precede Bio. JH is a consultant for or owns shares in the following companies: Artera, Kingmed Diagnostics, MoreHealth, OptraScan, York Biotechnology, Chimigen Bio, Sisu Pharma, ST N Technologies Limited and Sonablate Corp.

## Funding

This work was supported by the DoD Prostate Cancer Research Program (PCRP) Grant #HT94252310125 to Everardo Macias and the Early Investigator Research Award #HT94252510622 to Umar Mehraj. We thank Erik Soderblom, the Duke Proteomics Core, Department of Pathology, the Duke Cancer Institute Shared Resources, and the Cancer Center Isolation Facility, which are supported in part by the NCI Cancer Center Support Grant #P30CA014236. We thank Aurora Cabrera, Scott Lyons, Laura E. Herring, and Lee M. Graves (UNC Lineberger, Department of Pharmacology) and the UNC Proteomics Core Facility for assistance with MIB/MS experiments. The UNC Proteomics Core is supported in part by the NCI Center Core Support Grant (2P30CA016086-45) to the UNC Lineberger Comprehensive Cancer Center. The Structural Genomics Consortium (SGC) is a registered charity (No. 1097737) supported by funding from Bayer AG, Boehringer Ingelheim, Bristol Myers Squibb, Genentech, Genome Canada through the Ontario Genomics Institute [OGI196], the EU/EFPIA/OICR/McGill/KTH/Diamond Innovative Medicines Initiative 2 Joint Undertaking [EUbOPEN grant 875510], Janssen, Merck KGaA (also known as EMD in Canada and the US), Pfizer, and Takeda. Antonina Mitrofanova is supported by NIH NLM R01LM013236, ACS RSG-21-023-01-TBG, and DoD Data Science HT94252410346.

## Acknowledgements

We thank the Duke Proteomics Core, Department of Pathology, the Duke Cancer Institute Shared Resources, and the Cancer Center Isolation Facility, Duke University.

---

**Figure S1.**
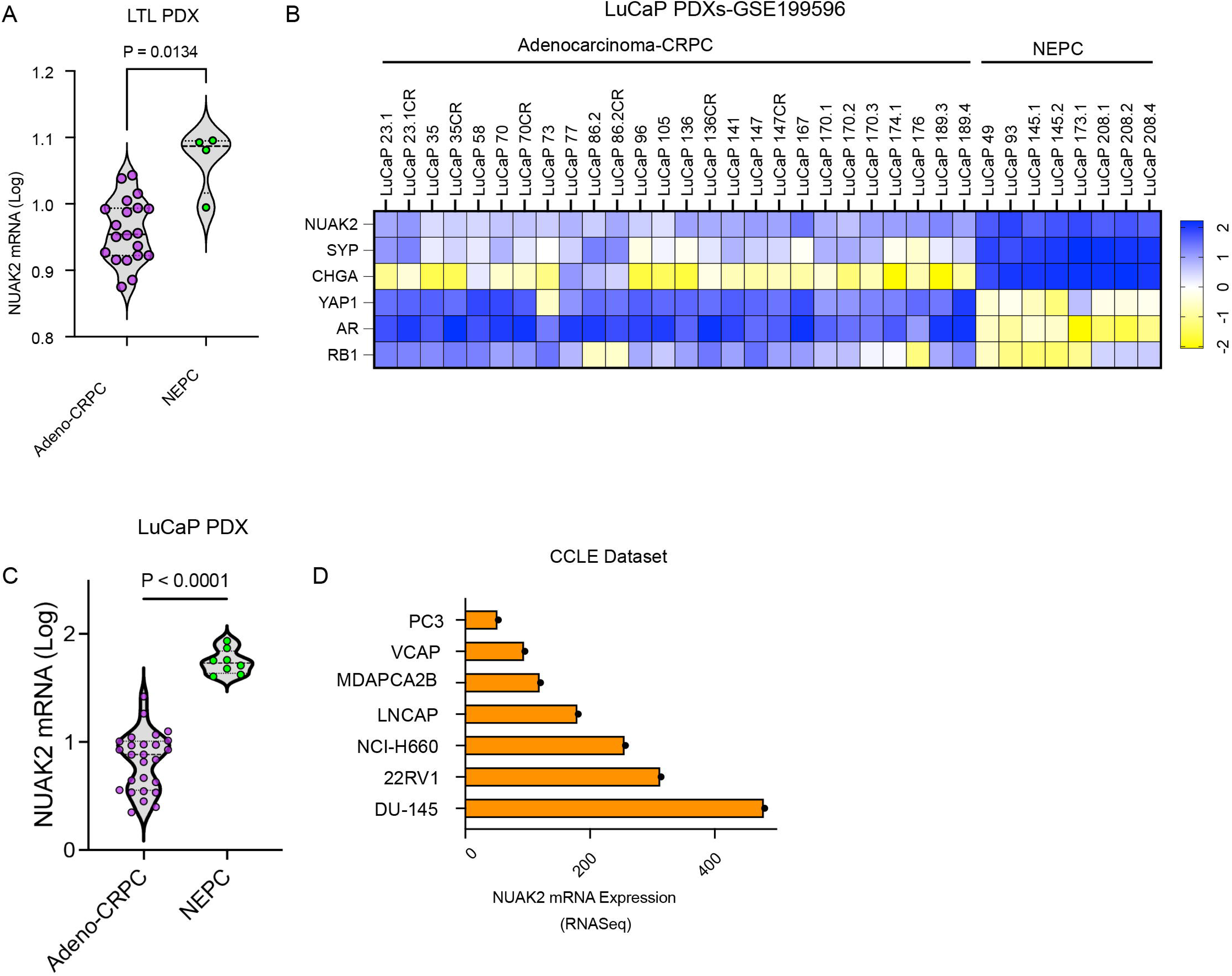
NUAK2 is upregulated in advanced prostate cancer. (A) Comparison of NUAK2 mRNA expression between adenocarcinoma-CRPC and NEPC LTL PDX models. (B) Heatmaps illustrating mRNA expression levels of denoted genes in adenocarcinoma-CRPC and NEPC LuCaP PDXs (30). (C) Comparison of NUAK2 mRNA expression levels between adenocarcinoma-CRPC and NEPC LuCaP PDXs (30). (D) NUAK2 expression profiles across PCa cell lines in the CCLE dataset. P values were determined by one-way ANOVA in (A) and unpaired two-tailed Student’s *t*-test for (C).

**Figure S2.**
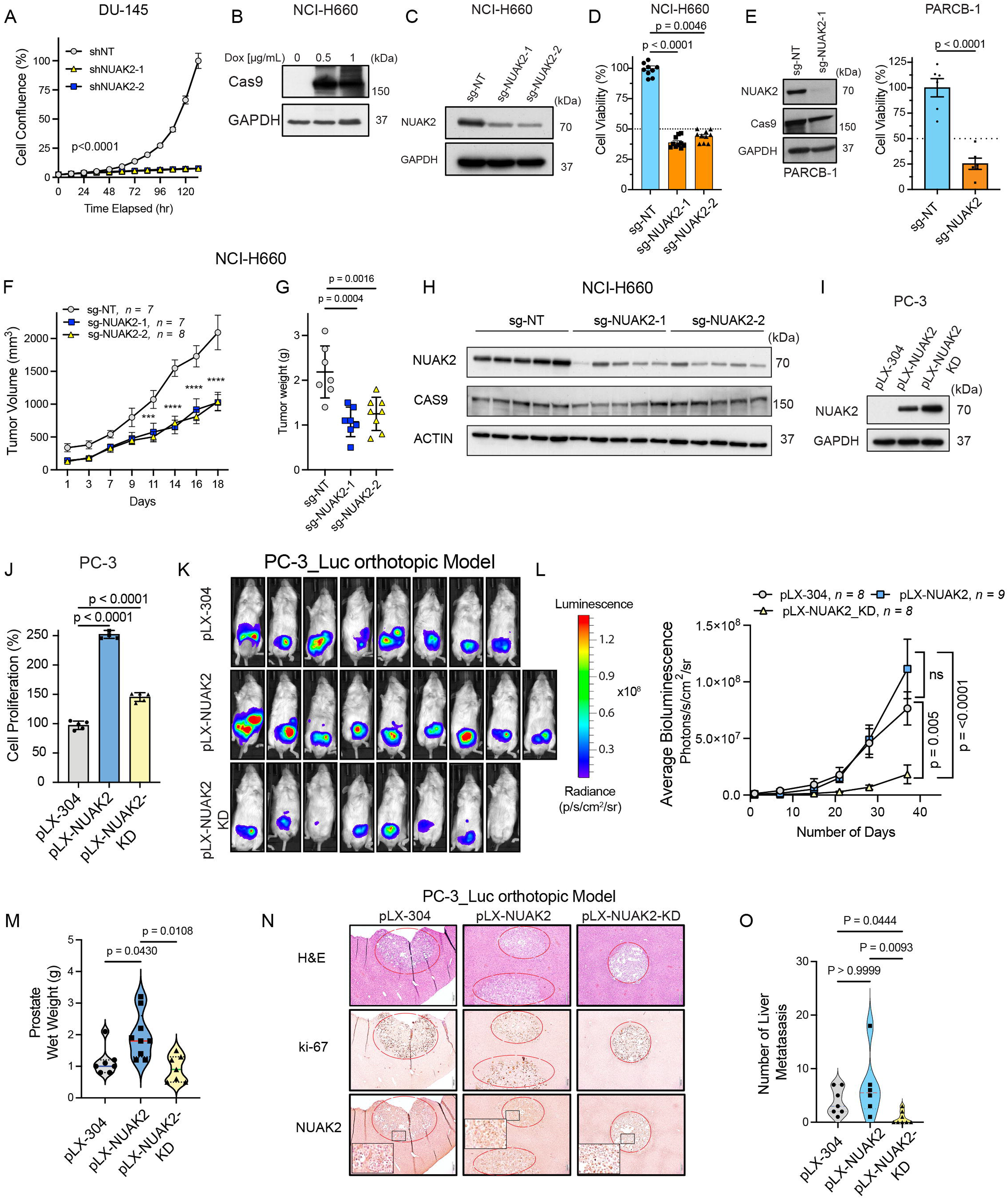
NUAK2 is essential for cell proliferation and tumor growth. (A) Cell confluence of DU-145 transduced with TRE3G shNT or shNUAK2 induced with doxycycline. Data are presented as mean ± SD. Statistical analyses were performed using two-way ANOVA. (B) Representative immunoblot analysis of Cas9 expression in TRE3G-Cas9-NCI-H660 cells treated with doxycycline for 72 h. (C) Representative immunoblot analysis of doxycycline-inducible NUAK2 knockout with two sgRNAs targeting NUAK2 in TRE3G-Cas9-NCI-H660 cells following doxycycline treatment (1 µg/mL) for 96 h. (D) Cell proliferation assay of NCI-H660 following doxycycline-induced NUAK2 knockout after 10 days. Data are presented as mean ± SD. P values were determined by one-way ANOVA. (E) Representative immunoblot validating doxycycline-induced NUAK2 knockout in PARCB-1 cells after 4 days and the corresponding cell proliferation assay after 10 days. Proliferation data are presented as mean ± SD. P values were determined using an unpaired two-tailed Student’s t-test. (F) TRE3G-Cas9-NCI-H660 xenograft tumor growth kinetics and (G) endpoint tumor weights from cells expressing inducible sgNT, sgNUAK2-1, or sgNUAK2-2 after doxycycline induction in male NSG mice. Data are shown as mean ± SEM; n = 7–8 mice per group. (H) Immunoblot analysis of individual TRE3G-Cas9-NCI-H660 xenograft tumors expressing sgNT, sgNUAK2-1, or sgNUAK2-2 constructs to confirm NUAK2 knockout and Cas9 expression *in vivo*. (I) Validation immunoblots of NUAK2 expression in PC-3 cells following ectopic expression of empty vector control (pLX-304), wild-type NUAK2 (pLX-NUAK2), or kinase-dead NUAK2 (pLX-NUAK2-KD). (J) Proliferation assay of PC-3 cells stably expressing pLX-304, pLX-NUAK2, or pLX-NUAK2-KD. Data are shown as mean ± SD. (K) Representative bioluminescence images and (L) quantification of primary tumor burden in PC-3_Luc cells expressing pLX-304, pLX-NUAK2, or pLX-NUAK2-KD. Data represent mean ± SEM, pLX-304 n = 8, pLX-NUAK2 n = 9, and pLX-NUAK2-KD n = 8. (M) Quantification of prostate wet weights at endpoint from PC-3_Luc tumor-bearing mice expressing pLX-304, pLX-NUAK2, or pLX-NUAK2-KD. pLX-304 n = 7, pLX-NUAK2 n = 9, and pLX-NUAK2-KD n = 7. (N) Representative hematoxylin and eosin (H&E), human-specific Ki67 and NUAK2 immunohistochemistry (IHC) images of metastatic tumor foci in mouse liver. (O) Quantification of metastatic tumor burden based on the number of metastatic lesions. pLX-304 n = 7, pLX-NUAK2 n = 6, and pLX-NUAK2-KD n = 8. Statistical analyses were performed using two-way ANOVA for panels (F and L), one-way ANOVA with Tukey’s multiple-comparison test (G and J) and Kruskal-Wallis test with Dunn’s multiple-comparison test for panels (M and O).

**Figure S3.**
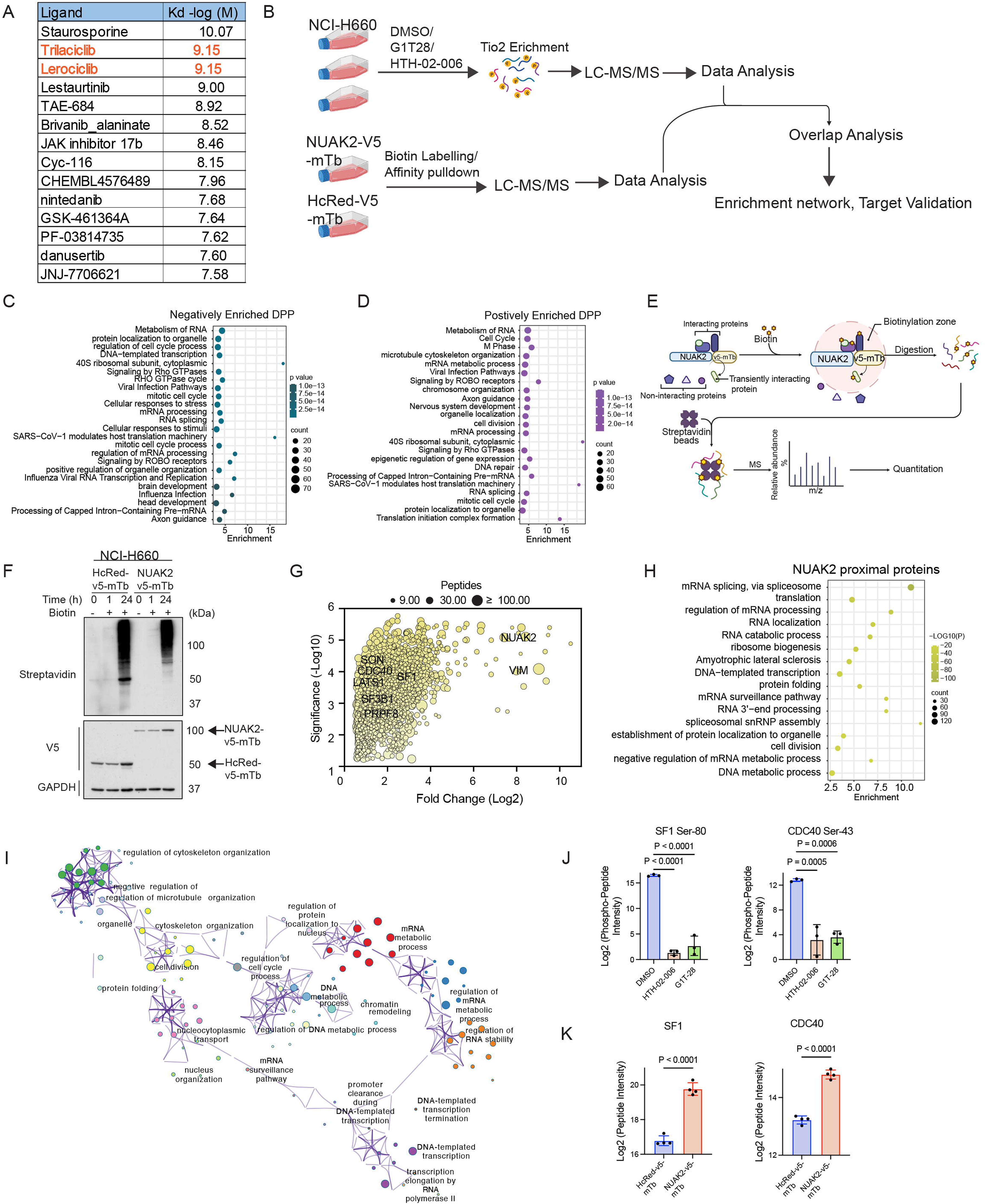
NUAK2 phosphoproteome and proximity networks implicate NUAK2 in RNA splicing. (A) Kinase inhibitors with documented NUAK2 binding activity identified through the Pharos database. (B) Schematic representation of the experimental workflow integrating label-free phosphoproteomic profiling and miniTurbo-based proximity labeling to characterize NUAK2-associated signaling networks in NEPC. Schematic was created with BioRender. (C–D) Pathway enrichment analysis of overlapping differentially phosphorylated peptides (DPP) with significantly altered phosphorylation (|log₂FC| ≥ 5) following treatment with either G1T-28 or HTH-02-006 relative to DMSO. (E) Schematic representation of the miniTurbo-based proximity labeling workflow. NCI-H660 cells stably expressing NUAK2-V5-mTb or HcRed-V5-mTb control were incubated with 0.5 mM biotin for 4 h, followed by streptavidin affinity purification of biotinylated proteins and identification by LC–MS/MS. (F) Immunoblot analysis of NCI-H660 cells stably expressing NUAK2-V5-mTb or HcRed-V5-mTb control, incubated with 0.5 mM biotin for 0, 1, and 24 h. (G) Bubble plot showing differentially biotinylated proteins by NUAK2-V5-mTb versus HcRed-V5-mTb (control). Each bubble represents a distinct protein, with the size of the bubble corresponding to the extent of biotinylation enrichment (number of peptides). (H) Enrichment analysis of proteins identified by NUAK2-dependent biotin labeling. Enrichment significance was determined by hypergeometric test; FDR < 0.05 considered significant. (I) Network visualization of enriched functional terms derived from clustered gene set enrichment analysis (GSEA) of 273 candidate proteins. Node colors indicate cluster identity. The network was rendered and visualized using Cytoscape v5. (J) Quantification of SF1 Ser80 and CDC40 Ser43 phosphorylation following pharmacological perturbation of NUAK2 with HTH-02-006 or G1T-28 in NCI-H660 cells, as determined by phosphoproteomic analysis. Data are presented as mean ± SD from three biological replicates. Statistical significance was determined by one-way ANOVA relative to vehicle control. (K) Quantification of SF1 and CDC40 enrichment following NUAK2 miniTurbo proximity labeling. Data are presented as mean ± SD from four biological replicates. Statistical significance was determined by unpaired two-tailed Student’s t-test comparing NUAK2-V5-mTb and HcRed-V5-mTb control samples.

**Figure S4.**
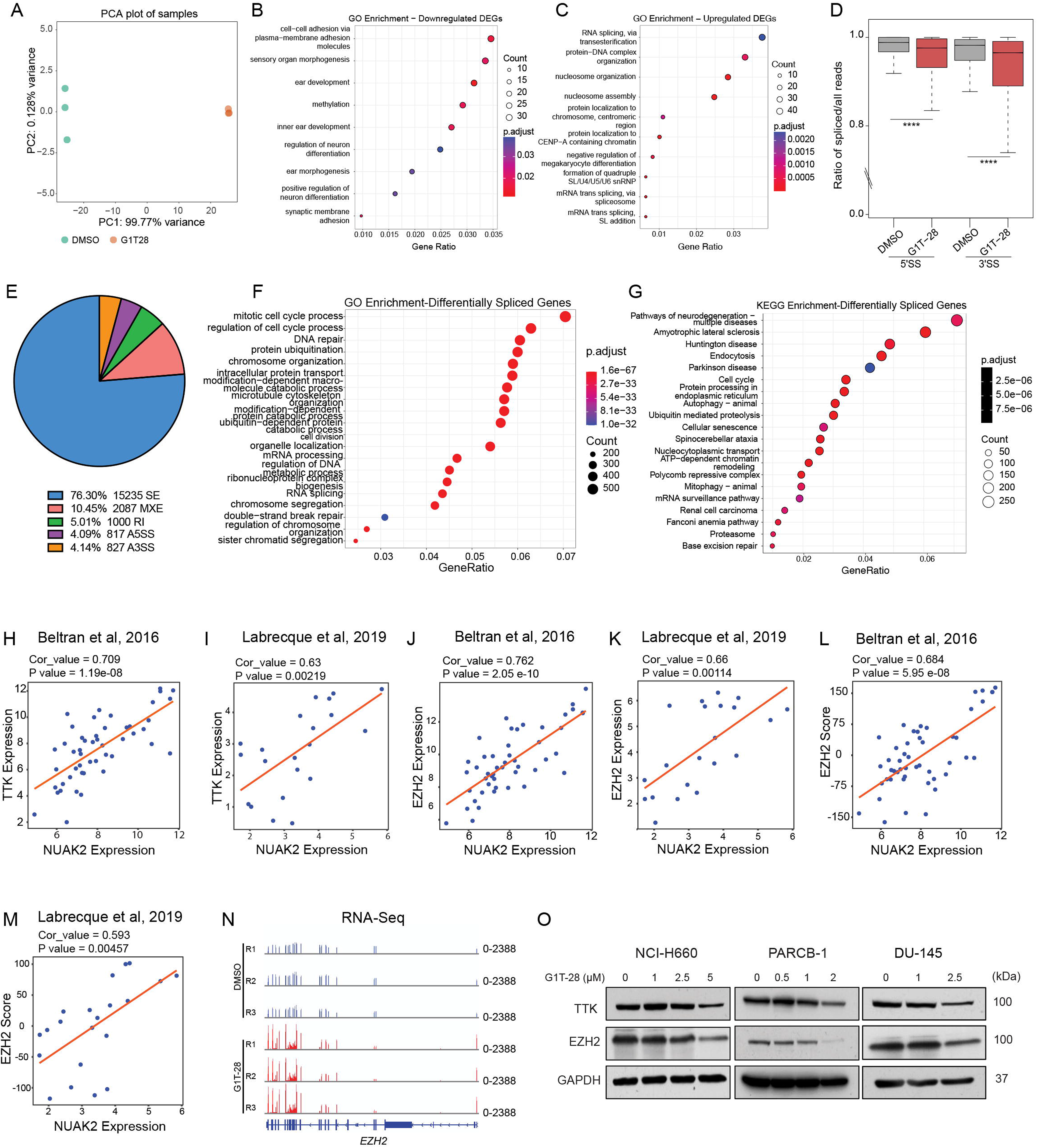
NUAK2 inhibition alters splicing landscapes. **(A)** Principal component analysis (PCA) of RNA-seq profiles showing clustering of NCI-H660 samples treated with G1T-28 or DMSO. (B–C) GSEA of differentially expressed genes in NCI-H660 cells following 24□h G1T-28 treatment. (D) Box plots depicting the ratio of spliced reads over total reads across all introns in 8,518 selected isoforms from RNA-seq data. SS denotes splice sites. Statistical significance was assessed by Wilcoxon signed-rank test (**** P < 2.2 × 10⁻¹□). (E) Summary pie chart of total differential splicing events identified by rMATS-turbo in NCI-H660 cells treated with DMSO versus G1T-28 after excluding events supported by <10 reads in either sample group and retaining events with |IncLevelDifference| ≥ 0.3. (F) GSEA of genes exhibiting splicing alterations upon G1T-28 treatment in NCI-H660 cells, as identified by rMATS. (G) KEGG enrichment analysis of genes exhibiting splicing alterations upon G1T-28 treatment in NCI-H660 cells, as identified by rMATS. (H–I) Correlation between NUAK2 and *TTK expression*, (J–K) *NUAK2 and EZH2* expression (z-scores) in the Beltran et al. 2016 (9), and Labrecque et al. 2019 (29) cohorts. (L–M) Correlation between EZH2 signature scores and *NUAK2* expression (z-scores) in the Beltran et al. 2016 (9), and Labrecque et al. 2019 (29) cohorts. (N) IGV tracks showing RNA-seq read coverage across the *EZH2* locus in NCI-H660 cells treated with G1T-28 or DMSO (control) for 24□h. (O) Immunoblot analysis of G1T-28-treated NCI-H660 (96 h), PARCB-1 (72 h), and DU-145 (72 h) cells.

**Figure S5.**
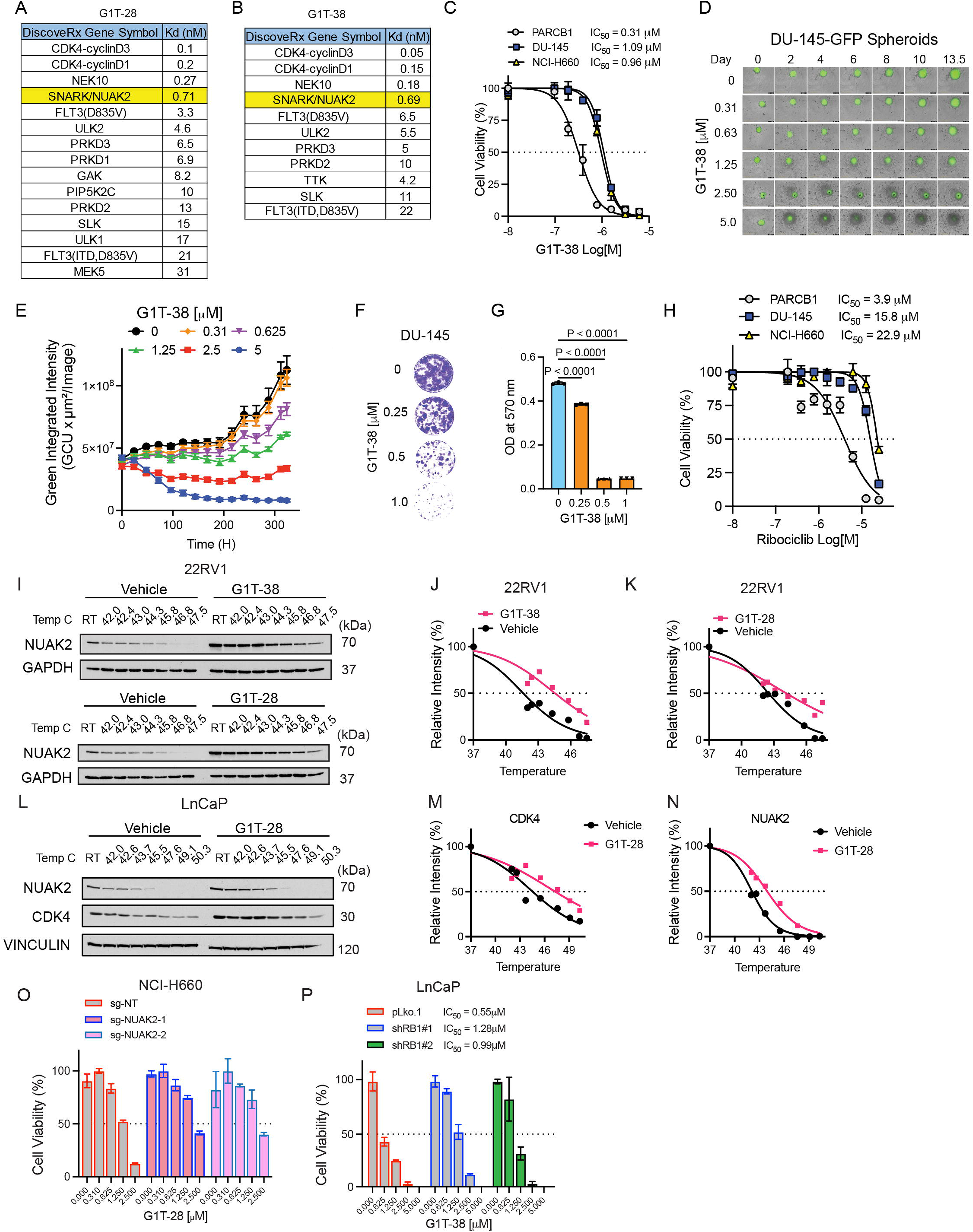
Kinase profiling and target engagement analyses of NUAK2-targeting compounds. (A) Kinases identified in kinome-wide screening of G1T-28 and (B) G1T-38 at 100 nM using the DiscoveRx platform, as reported by Bisi et al. 2016 and Bisi et al. 2017. (C) Dose response of NCI-H660, PARCB-1, and DU-145 following treatment with G1T-38. Data represent mean ± SD from three biological replicates. (D) Representative images of DU-145-GFP and (E) quantification of spheroid growth kinetics treated with G1T-38. Data represent mean ± SEM from three biological replicates. (F) Representative images and (G) quantification of DU-145 colony formation assay after G1T-38 treatment. Data represent mean ± SD from three biological replicates. (H) Ribociclib dose response curves of NCI-H660, PARCB-1, and DU-145 cells. Data represent mean ± SD from three biological replicates. (I) Representative CETSA immunoblot analysis and (J–K) corresponding NUAK2 melting curves of 22Rv1 cells treated with DMSO (vehicle), G1T-28 (6.25 µM), or G1T-38 (6.25 µM). (L–N) Representative CETSA immunoblot analysis (L) and corresponding CDK4 (M) and NUAK2 (N) melting curves in LNCaP cells treated with DMSO (vehicle) or G1T-28 (6.25 µM). (O) Cell viability analysis of NCI-H660 cells expressing non-targeting (sgNT) or NUAK2-targeting (sgNUAK2) sgRNAs following treatment with G1T-28 at the indicated concentrations for 8 days. Data represent mean ± SD from three biological replicates. (P) Cell viability of LNCaP cells expressing pLKO.1 control or independent RB1-targeting shRNAs (shRB1 #1 and shRB1 #2) following treatment with G1T-38 at the indicated concentrations for 96 h. Data represent mean ± SD from three biological replicates. IC_50_ and melting temperature were calculated using non-linear regression (curve fit) for panels (C, H, J, K, M, and P). Statistical analyses were performed using two-way ANOVA for panel (E) and one-way ANOVA with Tukey’s multiple-comparison test for panel (G).

**Figure S6.**
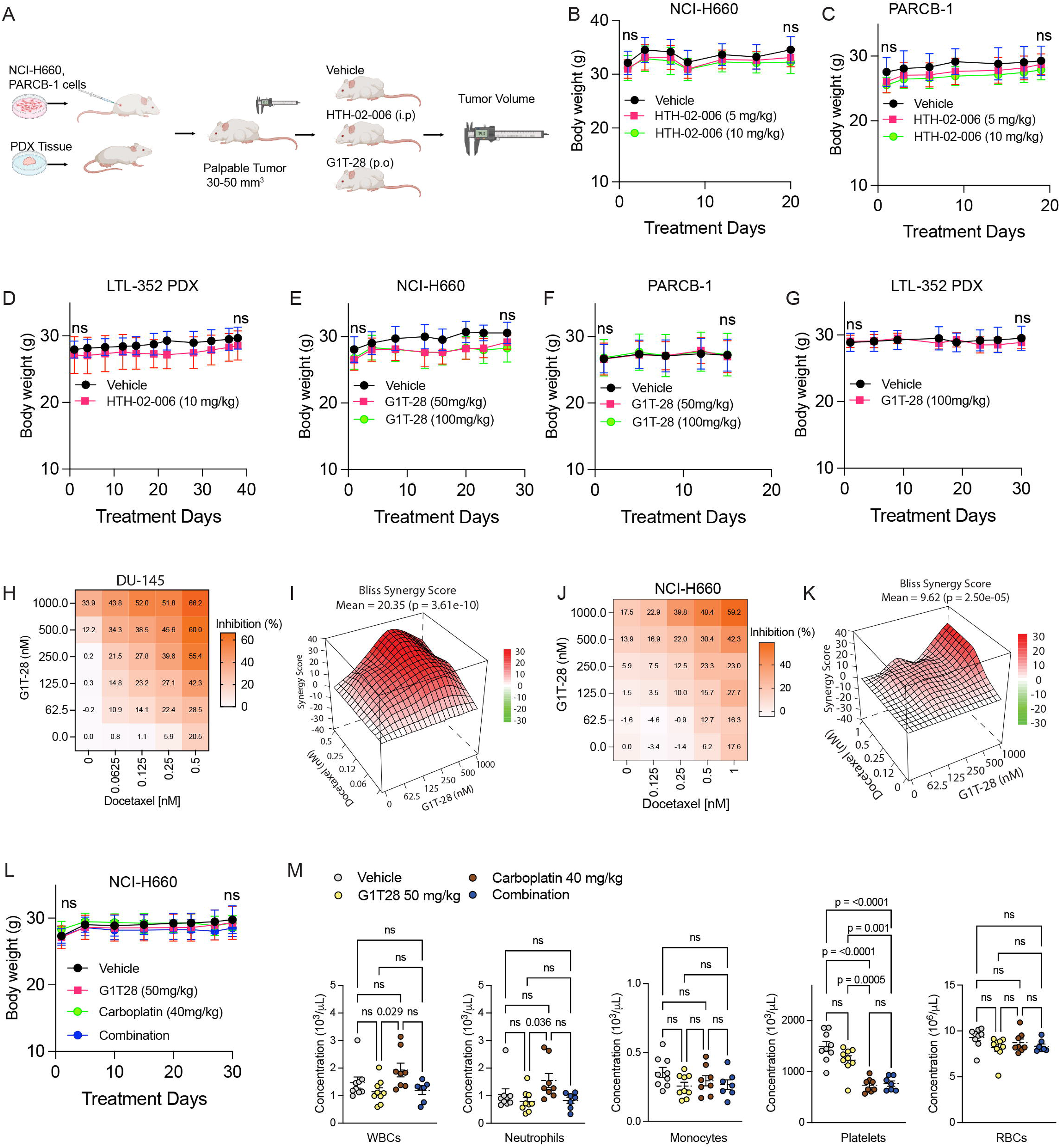
NUAK2 targeting synergizes with chemotherapy. (A) Schematic diagram illustrating the in vivo study design using NEPC cell line-derived xenograft (CDX) and NEPC patient-derived xenograft (PDX) models. Schematic generated using BioRender. (B–D) Average body weights of mice during treatment with vehicle or HTH-02-006 (5 or 10 mg/kg), and (E–G) vehicle or G1T-28 (50 or 100 mg/kg). Data are presented as mean ± SD. Statistical significance was assessed by two-way ANOVA. No significant differences in body weight were observed between vehicle- and treatment groups over the course of treatment at either dose (ns, not significant). (H) DU-145 G1T-28 and docetaxel dose combination matrix growth inhibition heatmap and (I) Bliss synergy plots. (J) NCI-H660 G1T-28 and docetaxel dose combination matrix growth inhibition heatmap and (K) Bliss synergy plots. (L) Average body weights of mice over the course of treatment with vehicle (control), G1T-28, carboplatin (40 mg/kg), or the combination. Data are presented as mean ± SD. Statistical significance was assessed by two-way ANOVA comparing treatment groups with vehicle control. No significant differences in body weight were observed among treatment groups (ns, not significant). (M) Complete blood count (CBC) analysis of mice following 26 days of treatment with vehicle, G1T-28, carboplatin, or the combination. Data are presented as mean ± SEM. Statistical analyses were performed using two-way ANOVA comparing vehicle and treatment groups.

## Notes

### Competing Interest Statement

AJ.A receives research support (to Duke) from the NIH/NCI, PCF, DOD, Astellas, Pfizer, Bayer, Janssen, BMS, AstraZeneca, Merck, Pathos, Amgen, Novartis. AJA declares consulting or advising relationships with Astellas, Pfizer, Bayer, Janssen, BMS, AstraZeneca, Merck, Exelixis, Novartis, Medscape, Telix, Duality Bio, MedIQ, IDEOlogy, Sumitomo, and Precede Bio. J.H is a consultant for or owns shares in the following companies: Artera, Kingmed Diagnostics, MoreHealth, OptraScan, York Biotechnology, Chimigen Bio, Sisu Pharma, ST N Technologies Limited and Sonablate Corp.

### Summary of Updates

The manuscript has been revised to incorporate additional experimental data, analyses, and textual revisions.

